# Cleaving DNA with DNA: cooperative tuning of structure and reactivity driven by copper ions

**DOI:** 10.1101/2023.06.06.543833

**Authors:** Sarath Chandra Dantu, Mahdi Khalil, Marc Bria, Christine Saint-Pierre, Didier Gasparutto, Giuseppe Sicoli

## Abstract

A copper-dependent self-cleaving DNA (DNAzyme or dexoxyribozyme) previously isolated by *in vitro* selection has been analyzed by a combination of Molecular Dynamics simulations and advanced EPR/ESR spectroscopy, providing insights on the structural and mechanistic features of the cleavage reaction at unprecedented resolution. The minimized 46-nucleotide deoxyribozyme forms duplex and triplex substructures that flank a highly conserved catalytic core. The self-cleaving construct forms a bimolecular complex that has a distinct substrate and enzyme domains. Cleavage of the substrate is directed at one of two adjacent nucleotides and proceeds *via* an oxidative cleavage mechanism that is unique to the position cleaved. The use of isotopologues of nucleotides allowed us to provide atomic resolution for the copper-substrate complex. The spectroscopic analysis overcomes the major drawbacks related to the ‘metal-soup’ scenario, also known as ‘super-stoichiometric’ ratios of cofactors *versus* substrate, conventionally required for the cleavage reaction within those nucleic acids-based enzymes. Our results pave the way for analysis on mixtures where metals/lanthanides are used as cofactors, having demonstrated that our approach may reach resolution of single nucleotide and beyond. Furthermore, the insertion of cleavage reaction within more complex architectures is now a realistic option towards the applicability of spectroscopic studies, both *in vitro* and *in vivo* matrices.

## Introduction/Main

DNA cleavage is a vital process in all living systems. For example, topoisomerase enzymes resolve topological problems of DNA in replication, transcription and other cellular transactions by cleaving one or both strands of the DNA.^1^ Another example are restriction enzymes (or restriction endonucleases), which protect the cell against virus infection by cleavage of the foreign DNA^2^ or by degrading cellular DNA during apoptosis of the affected cell.^3^ The activity of many anticancer drugs rely on their ability to introduce extended damage to the DNA in the (affected) cells (e.g. bleomycin),^4^ which can trigger apoptosis,^5^ leading to the cell death.^6^

DNA molecules, similar to RNA and proteins, are capable of folding into well-defined three-dimensional structures that can catalyze chemical reactions. Since the selection of the first DNAzyme,^7^ the number of DNA catalysts has dramatically increased and their catalytic repertoire has exceeded the cleavage of RNA.^8^ In addition, DNA-catalyzed reactions involving oligonucleotide substrates are DNA ligation,^9^ DNA cleavage,^10^ site-specific thymidine excision, DNA phosphodiester hydrolysis,^11^ DNA phosphorylation,^12^ DNA adenylylation/capping,^13^ and RNA ligation.^14^

The smallest self-cleaving domain that was obtained through *in vitro* selection was 69 nucleotides in length. The predicted secondary structure for this class of deoxyribozymes includes two stem-loops (I and II) that are interspersed with three single-stranded domains. In addition, a contiguous series of 21 nucleotides comprising the 31Z terminus of variant self-cleaving DNAs remained highly conserved throughout the *in vitro* selection process, indicating that these nucleotides are critical for deoxyribozyme function.^15^ By replacing the 26 nucleotides of the original 69mer DNA with a ‘tri-loop’, has been shown to enhance the stability of adjoining stem structures, a 46-nucleotide self-cleaving DNA (46mer) that retains this conserved sequence domain has been designed (Fig. 1a). The 46mer undergoes Cu^2+^-dependent self-cleavage with a rate that matches that of the original full-length deoxyribozyme.^16^

**Fig. 1.**
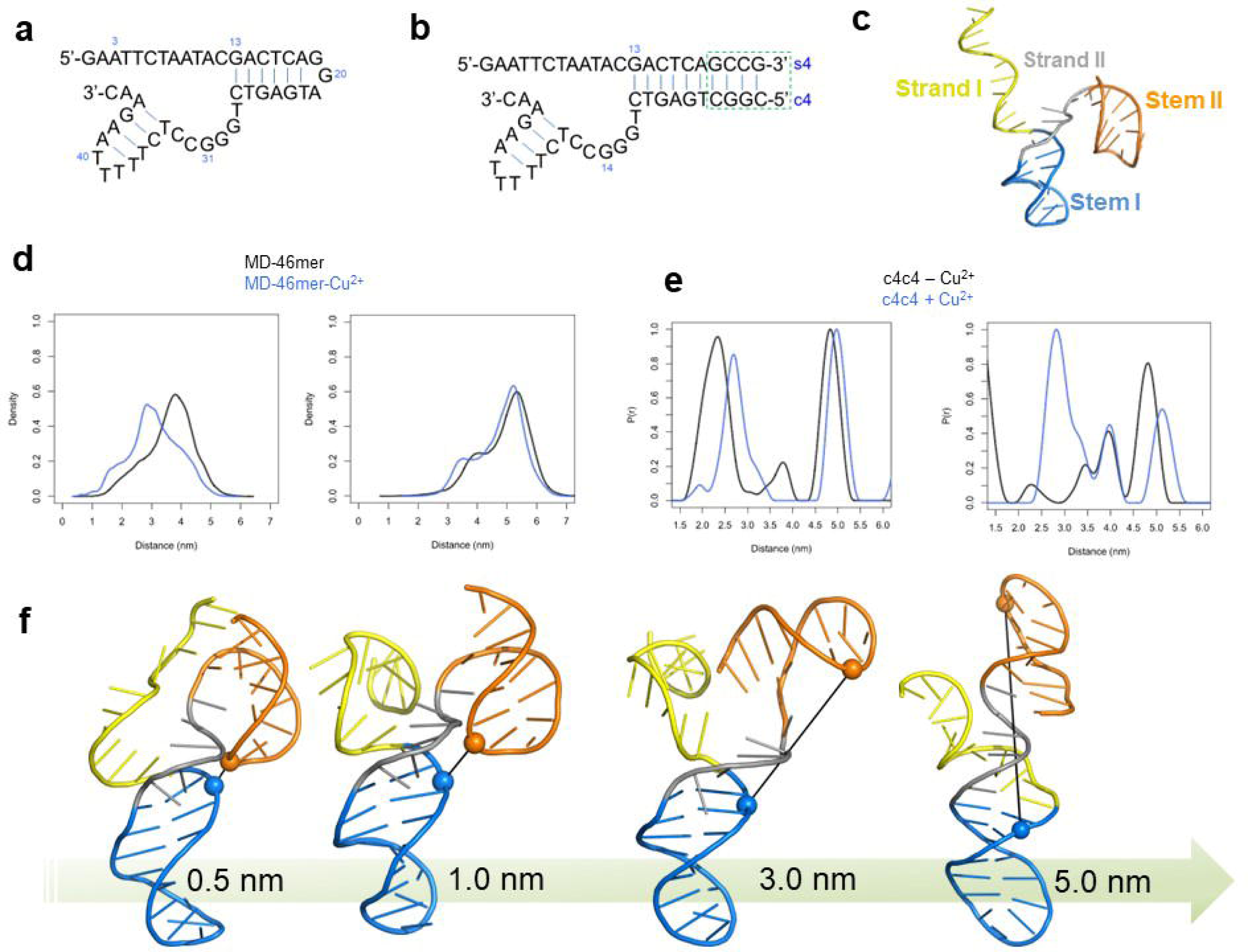
**a**, Secondary structures of the DNAzymes (deoxyribozymes) considered in this work. The 46mer as reduced structure containing the catalytic core. **b**, The bimolecular structure (c4s4) formed by substrate and enzyme oligomers, both containing four additional nucleotides (GCCG/CGGC base paring, c4s4 nomenclature deriving from those additional four bases replacing the GGA bulge fragment) to stabilize the secondary structure. **c**, Modelled structure of the 46mer for MD simulations; strand regions and stem regions are highlighted. **d**, Distance distributions profiles obtained from MD between the phosphate atoms of A14 and A41 and A18 and A41 exhibit conformational heterogeneity bon the absence and presence of Cu^2+^. **e**, Distance distributions on c4s4 structure derived by PELDOR/DEER experiments, recorded at t _0_ = 45 seconds (before cleavage); highly dynamic structure is confirmed by several populated states without and with Cu^2+^ cofactor. **f**, Structures from MD ensemble with Cu^2+^, selected based on distance between phosphate atoms of A14 (blue) and A41 (orange) ranging from 0.5nmto 5nm.

The 46mer can be divided into separate substrate and catalyst domains that are represented (Fig. 1b), for example, by the synthetic oligonucleotides s4 (substrate) and c4 (catalyst). The substrate DNA includes the primary site of DNA cleavage, while the catalyst DNA contains the original random-sequence domain that also contains all the nucleotides that were conserved during the *in vitro* selection process. The 22-nucleotide s4 and 29-nucleotide c4 DNAs do not undergo self-cleavage when tested individually under the permissive reaction conditions. However, when mixed, these DNAs form a bimolecular complex that retains full catalytic activity. Oxidation at the C11Z, C41Z, or C51Z positions of deoxyribose are putative sites for hosting the cleavage process.^17^ Random cleavage of DNA is readily observable upon incubation of polynucleotides in the presence of millimolar concentrations of mono-or divalent copper when combined with a similar concentration of ascorbate.^18^ Several DNA-cleaving agents such as 1,10-phenanthroline dramatically enhance the rate of DNA cleavage by Cu^2+^ and ascorbate, even when both Cu^2+^ and the DNA-cleaving agent are used at micromolar concentrations.^19^ Supplementing these oxidative cleavage reactions with H_2_O_2_ produces even greater rates of DNA chain cleavage, while enzymatically removing H_2_O_2_ from the reaction using catalase prevents the cleavage of DNA.^20^

Several significant drawbacks hinder the use of class II DNAs as artificial restriction enzymes for single-stranded DNA. First, the substrate strand contains nucleotides whose base identities are critical for catalytic activity. Therefore, the range of DNA sequences that can be cleaved is restricted thereby precluding the targeted cleavage of any DNA sequence. Second, the catalyst strand can also be cleaved during the reaction. Third, oxidative DNA cleavage yields some products that cannot easily be used in other molecular biology protocols due to the nucleoside fragments that are retained by some phosphate groups. Fourth, oxidative cleavage results in the loss of sequence information, as a base is destroyed during DNA strand scission. It is worth to note that a common denominator for the mentioned cleavage reactions is the ‘super-stoichiometric ratio’ (also known as ‘non-stoichiometric ratio’, or ‘metal-soup’ or ‘bath of electron spins’) used for the cofactor (i.e. Cu^2+^) which represents a deterrent for several conventional spectroscopic techniques. Thus, as *conditio sine qua non*, such an excess of cofactor is mandatory to realize the catalytic reaction under investigation and it shows severe limitations to the structural and mechanistic analysis of the chemical process.

RNA ligation catalyzed by DNAzyme may count on recent computational and crystal structures studies^21,22,23^, but these studies provide information exclusively on the ‘post-catalytic’ structure (product(s) of reaction) and not the pre-catalytic or catalytic complex. To date, insights on mechanistic and structural features related to DNAs-cleaved by DNAzymes, especially for catalytically active complex, are unknown. A recent study on RNA-cleaving DNA catalyst support the structural plasticity of the 10-23 DNAzymes architecture^8^; however, no paramagnetic species are involved as endogenous cofactor in such study, as most of the DNAzymes involve, and replacing the native cofactor with a paramagnetic species is then mandatory.

### 3D architecture of DNAzyme and cleavage reaction

Our integrative approach combining Molecular Dynamics simulations (MD) and advanced Electron Paramagnetic Resonance (EPR) spectroscopy (also known as Electron Spin Resonance, ESR) reveals information on the intrinsic structure of the DNAzyme and the DNAzyme in action (i.e. during the oxidative cleavage). Mass spectrometry, NMR-DOSY and MD techniques provide mechanistic and structural features of the global architecture of the DNAzyme, validating one of the putative process and selective sites of reactions. In Fig. 1a the 46-mer (as mono-molecular structure) is shown, and in Fig. 1b the corresponding bi-molecular structure (c4s4) is depicted. The additional four nucleotides added for the formation of the bi-molecular complex do not affect the cleavage reaction and it further stabilizes the 3D architecture formed by catalyst (c4) and substrate (s4). Putative base-paired architectures for the above mentioned DNAzyme are depicted in Supporting Information S1.

### Molecular dynamics for global structure and binding site(s)

A 3D model of the 46mer DNA has been modelled (Fig 1c) and has been used for MD simulations without (MD-46mer) and with Cu^2+^ (MD-46mer-Cu^2+^) (Supporting Information S2). The modeled secondary structure of the 46mer DNA (Fig. 1c), consists of two stem-loops (I and II) that are interspersed with three single-stranded domains. This secondary structure is very dynamic and yet stable in the MD simulations. Although stem I formation is important for DNAzyme function, the sequences that comprise this structural element can be varied as long as base complementation is retained (i.e. short DNA sequences have been tested in the past to reduce the size of the catalyst 69-mer and still keep the catalytic activity). The distance distributions using distances between phosphate atoms of selected residue pairs of Stem I and Stem II (A14-A41 and A18-A41) suggest a highly dynamic structure of the 46-mer (Fig. 1d) which is also evidenced from the EPR PELDOR/DEER data (Fig. 1e). In Fig. 1f the range of conformations accessible with Cu^2+^, from the MD ensemble of MD-46mer-Cu^2+^ are shown and an extended repertoire of 46-mer structures without and with Cu^2+^ are reported in Supporting Information S3 and S4. The presence of copper ion(s) shifts the distribution to the left for A14-A41; the conformational polymorphism of the 46mer is further evident from the distances measured between several pairs (C6-A21 and G19-C46) chosen to study the dynamic features of the 46-mer (Supporting information S5). To study the interaction of Cu2^+^ with the 46mer, contact between Cu2+ and the DNA nucleotides was analyzed with a distance cut-off of 0.4nm (Residence Time Analysis; RTA) and residence time was defined as the percentage of simulation time Cu^2+^ was within 0.4nm of the DNA (Fig. 2a and Supporting Information S6). Residues 5-10 and 27-34 and 40-46 had more interaction with Cu^2+^ (≥10% of simulation time) across the entire ten replicas of MD-46mer-Cu^2+^. In replica 1, the shortest distance between A14-A41 nucleotide pairs (0.5nm) for ∼20ns (Supporting Information S7 and Fig. 2b) can be observed; in this replica interaction between Cu^2+^ and nucleotides C6-A8, C12, G13, C27, C28, and G31-T34 for >10% of the simulation time (Fig. 2b-2c-2d) can be seen. The conformation with the shortest distance between A14 and A41 has a single Cu^2+^ ion that can simultaneously interact with the substrate and the catalytic sites from s4 and c4 strands, with a plausible pre-reactive complex geometry shown in Fig. 2d with interactions between Cu^2+^ and G13, A14, G29, A31, and A41.

**Fig. 2.**
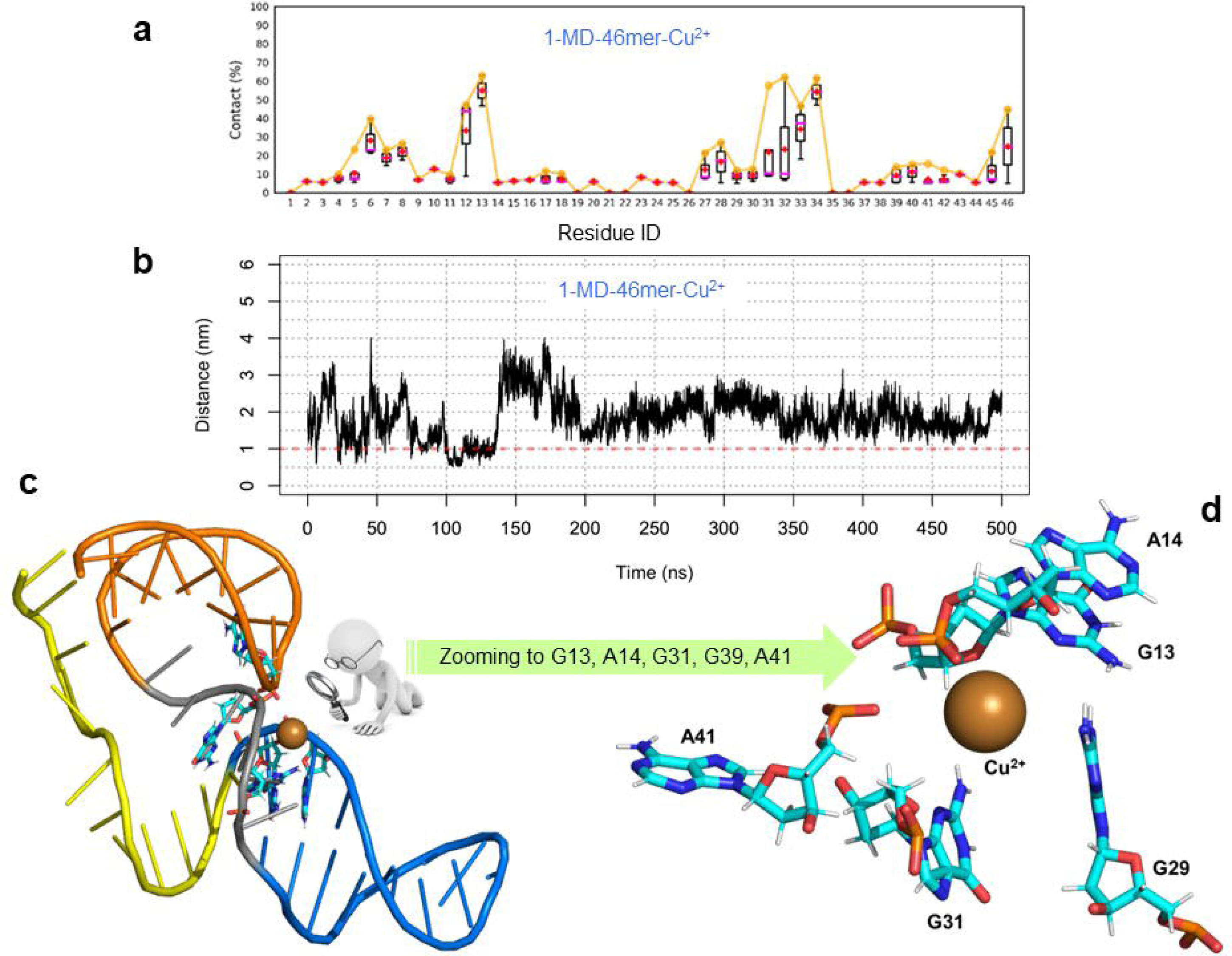
**a,** Interaction between Cu2^+^ and the 46mer with a distance cut-off of 0.4nm was defined as residence time and the contact frequency as percentage of simulation time is shown for 1-MD-46mer-Cu2^+^ **b**, Time series distance profile between the phosphate atoms of A14 and A41 from the MD trajectory of 46mer with Cu2^+^ (1-MD-46mer-Cu2^+^). **c**, Structure from 1-MD-46mer-Cu2^+^ with a distance of 0.5nm between A14 and A41, showing a plausible pre-reactive complex where **d**, Cu2^+^ is interacting with residues from substrate and catalytic strands simultaneously.

### EPR spectroscopy on c4s4 structure (bimolecular structure corresponding to **the 46-mer)**

The long-range distances evaluated by MD analysis have been compared (**Fig. 1e**) with the distance distributions obtained by PELDOR/DEER EPR techniques,^24-25^ used on the c4s4 architecture, labeled with two different spin probes (Supporting Information S8).^26-27^ The different populated states of c4s4 conformers detected by inter-spin distances agree with the scenario obtained by MD. The pronounced polymorphism exhibited by c4s4 without Cu^2+^ is not modified by the presence of the metal cofactor.

To go beyond the description of long-range interactions, short-range interactions between c4s4 and Cu^2+^ ions have been studied combining Continuous Wave (CW) and Pulse EPR/ESR spectroscopy (i.e. Hyperfine spectroscopy). As for copper chlorine in frozen solution and for monomeric ligand/Cu^2+^ complex previously described,^28^ spectral features typical for Cu^2+^ complexes with a (d_x_^2^__y_^2^)^1^ ground state (g⍰ > g ┴) are observed (Fig. 3a).^29^ The comparison of the g values and the Cu^2+^ hyperfine parameters between Cu^2+^ chlorine and c4s4 adduct indicates a metal–nucleobase interaction. The EPR parameters of Cu^2+^ chlorine are typical from Cu^2+^ complexes with four oxygen atoms as equatorial ligands.^29^ Instead the g values and Cu^2+^ hyperfine parameters of Cu^2+^/c4s4 complex are typical for Cu^2+^complexes with either three oxygen atoms and one nitrogen atom (Fig. 4b). Furthermore, the g values and the hyperfine parameters of samples containing c4s4 (all^14^N) and c4s4 (selectively labeled with^15^N) are very similar to each other (Fig. 4c). This suggests the presence of interactions between the Cu^2+^ ion and nucleobase(s).^30^

**Fig. 3.**
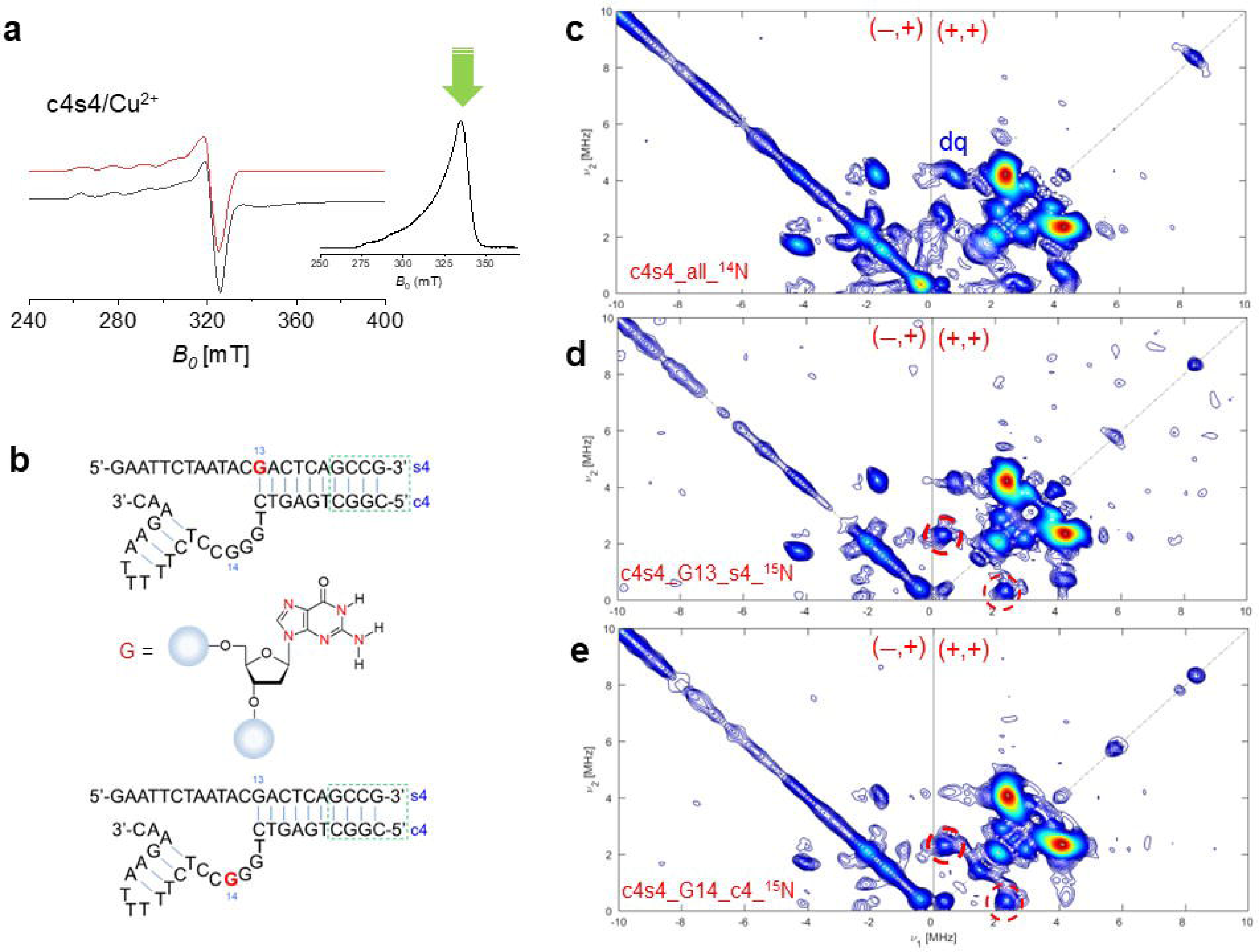
**a,** Continuous-Wave (CW, 9.3 GHz) EPR spectrum for c4s4/Cu^2+^ mixture(ratio 1:30), *black line*, and corresponding fit (*red line*). The Echo-detected Field-Swept (ED-FS, 9.72 GHz) spectrum where the observed field for pulsed experiments is indicated by the green arrow. **b**, DNA sequence selectively labeled with guanine (^15^N isotopically enriched) for the HYSCORE experiments. **c-d-e**, HYSCORE (HYperfine Sublevel COrRElation spectroscopy) 2D EPR experiments for the native c4s4 architecture. By comparing the (+/+) quadrant of the HYSCORE experiments for the unlabeled c4s4 with the samples containing guanosine G14 (c4) sequence and guanosine G13 (s4) additional cross peaks with ΔV 2.7 MHZ have been observed. These additional cross peaks are generated by the hyperfine coupling Cu^2+/15^ N (having^15^N nuclear spin I = 1/2, the manifold of the coupling generating a doublet without the presence of double quanta transition and/or frequencies combination).^15^N Larmor frequency (1.51 MHz) is indeed observed only on the HYSCORE experiments recorded on the sequences containing isotopologues (circled cross peaks. In addition, the comparison between the (―/+) quadrant of the unlabeled/labeled samples shows the reduced cross peaks (double-quanta transition for^14^N, indicated as dq).

**Fig. 4:**
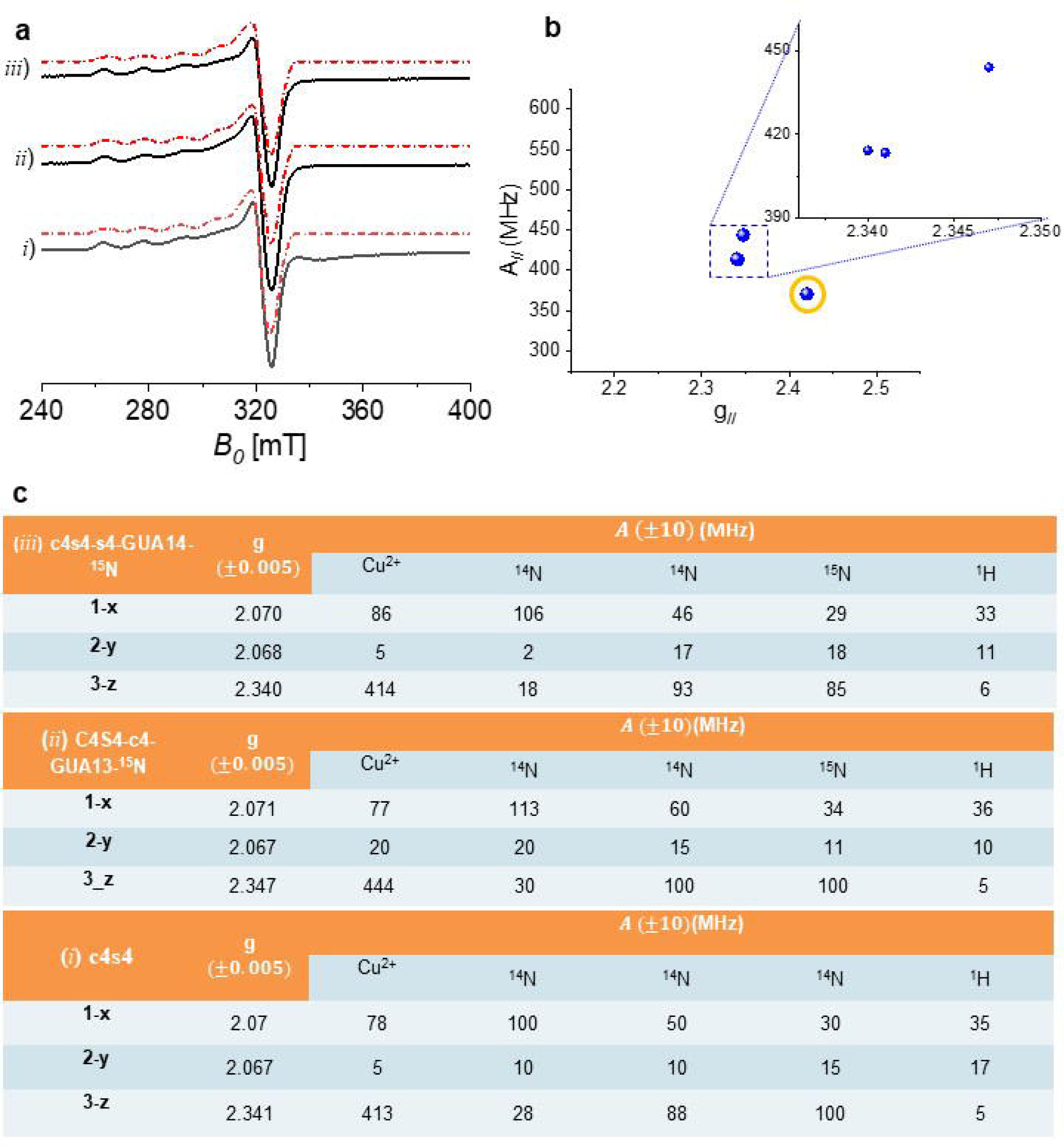
**a**, CW-EPR spectra for Cu-c4s4 sample at X-Band, measured at 120K. The red dotted lines are the fitting of the experimental spectra (black line). (*i*) c4s4 unlabeled sample, simulated using^63^Cu,^1^H and^14^ N, (*ii*) c4s4 isotopic labeled by^15^ N at Guanosine residue-13 of the substrate (s4); (*iii*) c4s4 isotopic labeled by^15^N at Guanosine residue-14 of the catalyst (c4). **b**, The Peisach-Blumberg plot; the hyperfine coupling with^63^Cu forthec4s4systemshasbeenplotted(asfunctionofg _//_, including the references (CuSO _4_, circled blue point). **c**, Table summarizing the hyperfine couplings with^63^Cu,^14^N,^1^H and^15^N.

Although the CW EPR data provide evidence of a coordination of the Cu^2+^ metal by c4s4 systems, detailed information about the metal ion site cannot be obtained by 1D-CW experiments. Thus, further insights into the metal ion site is provided by 2D-pulsed EPR techniques. To get information about the weakly coupled nuclei, the 2D-ESEEM (known as HYSCORE) technique has been used. In Fig. 3a inset, the echo-field swept spectrum and the observed field (green arrow) chosen for the HYSCORE experiment. Furthermore, the hyperfine spectroscopy has been supported by the use of isotopologues on selected residue (i.e. nucleotides, Fig. 3b). The choice of isotopically enriched residues has been driven by the RTA analysis from MD, in order to exploit the potentiality of HYSCORE, despite the excess of Cu^2+^ cofactor. The HYSCORE 2D-spectrum of c4s4 bimolecular complex formed with Cu^2+^ (Fig 3c) confirms *a plethora* of weak coupling with^14^N nuclei within the base-pair arrangement. Based on the RTA from MD simulations,^15^N isotopic labeled nucleobases, G13 on the substrate and G14 on the catalyst, have been used to ‘switch on’ the selective coupling suggested in the previous MD section.

The HYSCORE spectrum of the Cu^2+^/c4s4 complex also shows two types of weakly coupled protons. One type is characterized by an intense ridge close to the antidiagonal at the proton Larmor frequency (14 MHz) and can be assigned either to protons of water molecules coordinated in axial positions or/and to weakly coupled protons of solvent molecules.^29^ A more crowded scenario is reserved to the^14^N hyperfine coupling spectral regions; as the MD analysis has suggested, several sites are coordinated to Cu^2+^, involving several^14^N nuclei. The low-frequency regions of the HYSCORE spectra of samples c4s4 and c4s4 (G13-s4-^15^N, G14-c4-^15^N) are reported in Figure 3c-e. The HYSCORE spectrum of Cu^2+^/c4s4 (all-^14^ N) is dominated by cross-peaks that are assigned to double-quantum (DQ) correlation peaks from^14^ N. Furthermore base-pairing around the copper coordination increases the number of^14^ N nuclei, as it can be observed in the HYSCORE experiments. As previously reported for guanosine mono-phosphate,^29^ also quadrupole interaction can be considered (|e^2^qQ/h| = 3.02 ± 0.05 MHz) (Supporting Information S9, HYSCORE fitting at X-band, Table S10). Even if those samples contain an excess of cofactor (Cu^2+^) the doublet/cross peaks for^15^N is successfully achieved. Indeed, the assignment to the remote nitrogen is reinforced by the analysis of c4s4 containing isotopologues (^15^ N) and a similar remote nitrogen pattern is obtained in addition to cross-peaks derived by interaction with^14^N nuclei. Furthermore, within the base-pair arrangement, it possible to discard N3 and N9 of guanine within the coordination sphere; those are involved within the hydrogen bonding network of the minor groove of DNA duplex form and oriented far from the metal center.

The HYSCORE spectrum at 9.7 GHz (X-band) is characterized by single-quantum, double quantum, and combination peaks of nitrogen nuclei. Cross-peaks related to^15^N are observed for the G14 on the catalyst (c4 for the complex c4s4, numbered at G31 on the 46-mer) and on G13 (substrate). To discard any ambiguities, HYSCORE experiments have been recorded at higher frequency/field (34 GHz, Q-band). Those experiments confirmed the coupling of Cu^2+^ with^15^ N on the guanine of the catalyst with higher resolution because of cancellation conditions for different nitrogen nuclei (Fig. 5a and 5b).

**Fig. 5:**
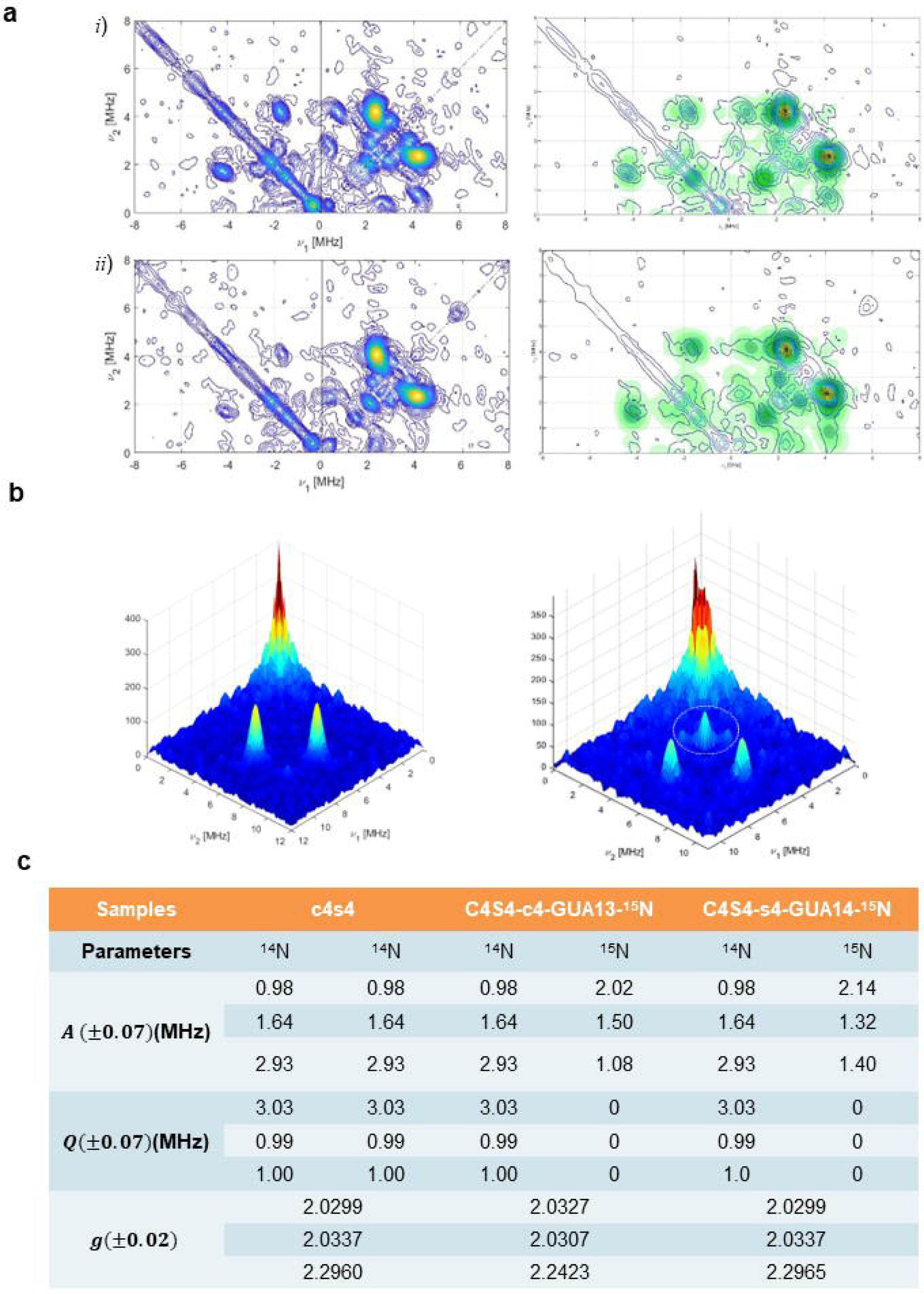
**a**, HYSCORE spectra recorded at X-band (9.7 GHz) (*left*) and corresponding fit (*right*) for (*i*) c4s4 unlabeled sample, simulated using^63^Cu,^1^H and^14^N, (*ii*) c4s4 isotopic labeled by^15^N at Guanosine residue-13 of the substrate (s4); **b**, 3D plot for the HYSCORE spectra recorded at Q-band (34 GHz) for (*i*) c4s4 unlabeled sample, simulated using^63^ Cu,^1^H and^14^ N, (*ii*) c4s4 isotopic labeled by^15^N at Guanosine residue-13 of the substrate (s4); **b**, Main parameters used for HYSCORE fitting procedure using^63^Cu,^14^N,^1^H and^15^N.

### Monitoring the cleavage reaction by MALDI-TOF experiments and NMR-DOSY experiments

Besides the local interactions deciphered by combining MD analysis and EPR/ESR 1D and 2D techniques, mass spectrometry and NMR diffusion-ordered spectroscopy (DOSY) allowed us to validate the products of the cleavage reaction as well as radical intermediates. Hydroxyl radical generated by a Fenton-like reaction^31^ has been detected by spin-trap technique (Fig. 6a and Supporting Information S11); the products of cleavage reaction (3’-phosphoglycolate and 5’-phosphate fragments, Fig. 4b) confirmed by NMR diffusion (DOSY) (Table S13 and Supporting Information S12-S17) and MALDI-TOF experiments (Fig. 6c-d), able to discard putative hydrogen abstraction (Supporting Information S18-S28).

**Fig. 6.**
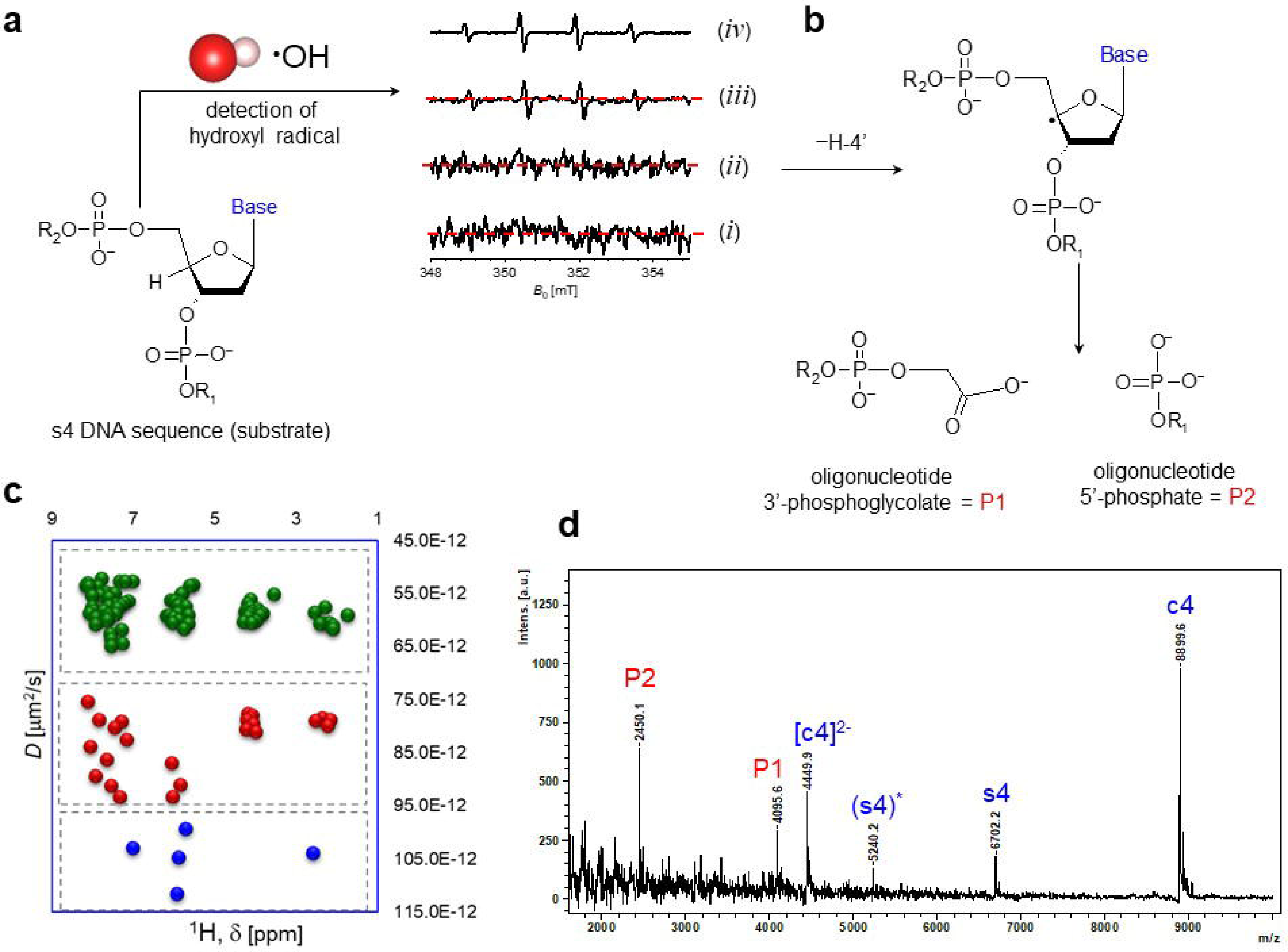
**a**, EPR spin-trap experiments on c4s4, confirming the generation on the HO• [DMPO-OHadduct]•radicalspecies(asforaFenton-typereaction);thespin-trap experiments carried out on the buffer solution without (*i*) and with DMPO (*ii*) do not show the production of hydroxyl radical, while in the presence of Cu^2+^ (*iii*) and Cu^2+^/c4s4 (*iv*), the typical signal of hydroxyl radical is observed. **b**, Proton abstraction from C4’ position on the ribose ring (under aerobic conditions) is here reported in una simplified scheme (selected steps are omitted). The product 3’-phosphoglycolate (P1) and the 5’-phosphate (P2) are the two DNA fragments derived by the selective cleavage. **c-d**, Diffusion Ordered SpectroscopY (DOSY) experiments and Mass Spectrometry validate the formation of P1 and P2; the diffusion coefficients have been correlated to the molecular weight (MW) of the different fragments, while the MALDI-TOF experiments, direct detection of the fragments upon cleavage reaction has been obtained.

## Discussion

The snapshot from the MD simulation of MD-46mer-Cu^2+^ from replica 1, at 104.65ns with distance of 0.53nm between A14 and A41, (Fig. 2b) suggests the assignment to a ‘pre-reaction complex’ (Fig. 2c), with a single Cu^2+^ interacting with both the substrate, G13, and the catalytic arm G31 simultaneously. Such structure gives insights into the specificity of the binding, as confirmed by the EPR data. If EPR hyperfine spectroscopy has been established as method of choice for Fe-S cluster proteins,^32^ mechanistic insights into DNAzymes are strongly affected by multiple binding sites.^24^ The combined approach (EPR/MD/NMR/MALDI) can focus on the selected guanine residue(s) with unprecedented resolution. Furthermore, the role of the copper ion(s) has been deciphered both as structural (for the coordination on the catalytic core) and as mechanistic (for the oxidative step)^33^ as far as hydrogen abstraction on the C4’ is involved (as confirmed by the products of the cleavage). Such a cooperative role of the copper ion identified by EPR/ESR hyperfine spectroscopy is in agreement with a fitting of EPR/ESR binding isotherms recently proposed for the manganese ions^8, 34^ and with the few cooperatively binding modes proposed for the DNAzymes.^35-38^ The cooperative binding is combined with the intrinsic flexibility of the c4s4 (46mer) architecture.^39-40^

## Conclusions

Artificial enzymes based on DNA strands have a number of characteristics, such as chemical stability and ease of synthesis, that make them well suited for a number of practical applications. Attractive features for a next generation and more useful DNA-cleaving DNA enzyme are user-defined substrate specificity with minimal demands on the sequence of the target DNA, rapid and multiple turn-over kinetics under mild reaction conditions, and a small size to allow efficient large-scale chemical synthesis. The spectroscopic protocol here proposed combined with MD analysis could point out both mechanistic and structural features unknown to date. It paves the way for increasing the complexity both of cofactors (i.e. metal and lanthanides) and of structural features (i.e. analysis of DNAzymes containing several branched regions). Ligation reaction of RNA and/or DNA strands is the next suitable candidate for the approach presented in this work.

## Supporting information

Supplementary information list

## Acknowledgements

For the EPR experiments financial support from the IR INFRANALYTICS FR2054 is gratefully acknowledged. The authors thanks H. Ahouari (Chevreul Institute, Villeneuve d’Ascq) for technical assistance in recording CW spectra and troubleshooting. Prof. C. Hobartner (University of Wuerzburg) is kindly acknowledged for providing the cytosine TEMPO-labeled oligomers. MD simulations were performed using the HPC time granted via the UK High-End Computing Consortium for Biomolecular Simulation, HECBioSim 2(http://hecbiosim.ac.uk), supported by EPSRC (grant no. EP/R029407/1).

## Author contributions

S. C. D. performed the MD analysis; C. S.-P. et D.G. recorded the MALDI-TOF spectra and perform the analysis; M.B. recorded and analyzed the NMR experiments. M.K. analyzed the HYSCORE spectra and carried out the fitting of 1D and 2D experiments. G.S., S.C.D. conceived the EPR/ESR experiments and MD tailored approach; G.S., S.C.D. wrote the manuscript. All authors commented on the manuscript.

## Additional Information

The authors declare no conflicts of interest.

## METHODS

### Sample preparation for EPR experiments

Duplexes have been formed by heating DNA strands at 90°C for 3 minutes and slowly cooled down to room temperature. For the different pH, three buffers were used: Sodium Acetate (pH 4.00), Cacodylate (pH 7.45), Glycine/NaOH (pH 10.3). 10% (v/v) of glycerol was added before freezing the sample in liquid Nitrogen. With respect to Cu^2+^ solution (CuCl^2^), a molar excess of monomeric ligand or double helix was used, up to an excess of 1:10. 100 mL solutions were used in 4 mm tubes (X-band) and 3 mm tubes (Q-band), respectively.

### EPR experiments

Continuous Wave (CW) X-Band measurements were carried out using an X-band Bruker E500 instrument (9.4 GHz, TE^012^ resonator) equipped with a nitrogen flow cryostat. All CW experiments were recorded at 120 K and with a shot repetition rate of 100 Hz, unless stated otherwise. Pulsed EPR experiments at X-band were performed on a Bruker ELEXYS E-580 X-band spectrometer with a SuperX-FT microwave bridge and a Bruker ER EN4118X-MD4 dielectric resonator. Cryogenic temperatures (20 K) were obtained by the use of an Oxford flow cryostat. The field-swept EPR spectra were recorded by electron spin echo (ESE) detection; electron-spin-echo (ESE)-detected EPR experiments were carried out with the pulse sequence: π/2‒τ‒π‒τ‒echo. For the X-band experiments the mw pulse lengths t _π/2_ = 16 ns and t _π_ = 32 ns and a τ value of 200 ns were used. A two-step phase-cycle was applied to remove all unwanted echoes. The Hyperfine Sublevel Correlation (HYSCORE) experiments were carried out using the pulse sequence π/2‒τ‒π/2‒t _1_‒π‒t_2_‒π/2‒τ‒echo. The time traces of the HYSCORE spectra were baseline corrected using a third-order polynomial, apodized with a Hamming window and zero-filled. After two-dimensional Fourier transformation, the absolute value spectra were calculated. A four-step phase cycle (for X-band experiments) was used to remove unwanted echoes. The pulse sequence for the four-pulse DEER experiment was π/2 _obs_1Z−1Z*τ* _1_1Z−1Zπ_obs_1Z−1Z*t*_1_1Z−1Zπ_pump_1Z−1Z(*τ* _1_1Z+1Z*τ* _2_1Z−1Z*t*_1_)1Z−1Zπ_obs_1Z−1Z*τ* _2_. The pump pulse was applied on the spectral maximum and the observer pulses were applied at a frequency offset of 551ZMHz. Measurements on the Bruker Elexsys E580 spectrometer were acquired with a pulse delay *τ* _1_ of 2001Zns and a dead time delay of 1001Zns. All DEER data were analyzed using DeerAnalysis2022 based on MATLAB. The distance distribution *P*^dis^(*r*) was fitted by Tikhonov regularization (using residual method for regularization parameter selection). The capture of active radicals generated during the reaction was examined by EPR spectra (Bruker ELEXSYS 500 spectrometer) using DMPO as a spin trapping agent (Supplementary methods). DMPO (0.8 M) was immediately added to the c4s4 solution and transferred to a glass capillary tube. Then, the capillary tube was placed into a quartz EPR tube and EPR spectra were recorded. Typical spectrometer parameters are shown as follows, scan range: 100 G; center field set: 3510 G; time constant: 1.25 ms; scan time: 40.96 s; modulation amplitude: 0.5 G; modulation frequency: 100 kHz; receiver gain: 1.00 × 103; microwave power: 19.17 mW. Signal fitting was carried out using the Spin Fit program (Bruker). Signal fitting for HYSCORE experiments has been carried out using the HYSCOREAN software (https://epr.ethz.ch/software.html).

### Modelling and MD simulations

The 3D structure of the 46-mer (Fig. 1a) was modelled using x3dna package (v2)^41^ using 1JVE pdb^42^ as template for the “GGA hairpin”. Parmbsc1 forcefield^43^ was used for the MD simulation of the 46mer using gromacs2016 package^44^ 46-mer was placed in a dodecahedron box with 1.0 nm distance between the box walls and the DNA molecular and was solvated with tip3p^45^ water molecules (MD-46mer). For simulations with Cu2+ ions, 10 mM of Cu2+ ions were randomly placed in the simulation box (MD-46mer-Cu^2+^) (Fig. S2). Simulation system was energy minimized using steepest descent algorithm until the largest force was smaller than 1000 kJ/mol/nm, followed by temperature equilibration to 300 K in 100ps using Berendsen thermostat with a tau-t of 0.2 ps. Pressure was equilibrated to 1 atm in 1 ns using Berendsen barostat and temperature was regulated using Berendsen thermostat at 300 K.^46^ Production run simulations were started using the equilibrated structures and ten replicas were simulated for 500 ns. For production run simulations, temperature was regulated usi velocity-rescaling thermostat^47^ and pressure with Parrinello-Rahman barostat^48^ at 300 K and 1 atm using tau-t of 0.1ps and tau-p of 2ps. Structures were saved every 10 ps. Clustering of the MD simulations was done using gromos algorithm^49^ with snapshots sampled at 200 ps intervals using only phosphate backbone atoms with a RMSD cut-off of 0.2 nm. All the analysis was done using in-house python scripts and data visualization was done using Pymol.^50^

### NMR experiments

NMR experiments^51-56^ were recorded with a 3-mm NMR sample tube at 298°K using the following Bruker spectrometers with *z*-axis pulsed field gradients: Avance NEO at 900-MHz (21 Tesla) proton frequency with CPTCI CryoProbe is a proton-optimized triple resonance NMR ‘inverse’ probe. A standard pulse sequence lebpgp2s from Topspin 4.1, (Bruker) was used for diffusion experiments on the 900-MHz (21 Tesla) instrument. In total, 65 536 points with 16 scans were recorded in the proton dimension for each one dimension with variable diffusion gradient strength ranging between 2 and 95% in various steps. The following parameters were used: diffusion time (*Δ*) 0.0851Zs, gradient pulse (*δ*) 2500 µs smoothed rectangular-shaped gradients SMSQ10.100, relaxation delay (d1) 101Zs. NMR experiments were recorded with a 5-mm NMR sample tube at 298°K Avance NEO at 4001ZMHz (9.4Tesla) equipped withaTBIprobe.AllspectrawereprocessedwithTopspin4.1(Bruker)andDOSY treatment Software Dynamic center (Bruker)

### MALDI-TOF experiment

The MALDI-TOF^57-59^ mass spectra measurements were performed in the negative mode on a Microflex mass spectrometer (Bruker, Wissembourg, France). Prior to mass analysis, oligonucleotides solutions were purified and concentrated using Zip Tip pipette tips (Merck Millipore) filled with 0.6 µL C18 resin. A mixture of the purified DNA sample (10 pmol, 1 µL) was added to the matrix (3-hydroxypicolinic acid in 10 mM ammonium citrate buffer) and spotted on a polished stainless target plate using the dried droplet method. Spectra were calibrated using reference oligonucleotides of known masses. Each spectrum were obtained by summing 300 shots by the use of a 337 nm pulsed nitrogen laser beam (60 Hz). Linear mode was run with optimized voltages for ion sources (IonSource-1: 20 kV, IonSource-2: 18.5 kV) and pulsed ion extraction delay was fixed at 100 ns. In order to eliminate the intense low masses of the spectra (matrix peaks, solvents clusters) which normally saturates microchannel plate detectors, and with the aim to enhance the ratio signal to noise, all ions with less mass than 1800 Daltons were deflected. Spectra were accumulated by FlexControl Software (v.3.3.108.0) and processed with FlexAnalysis using Savitsky-Golay algorithm (with 0.2 m/z, one cycle) and baseline subtraction (Top Hat).

## References

1. Sangeetha Gowda, K. R., Blessy, B. M., Sudhamani, C. N., Bhoijya, H.S. Mechanism of DNA binding and cleavage Biomedicine and Biotechnology 2, 1–9 (2014).

2. Yuan, R., Bickle, T.A., Ebbers, W., Brack, C. Multiple steps in DNA recognition by restriction endonuclease from E. coli K Nature 256, 556–560 (1975).

3. Nagata, S., Nagase, H., Kawane, K., Mukae, N., Fukuyama, H. Degradation of chromosomal DNA during apoptosis Cell Death & Differentiation 10, 108–116 (2003).

4. Chen, J., Ghorai, M. K., Kenney, G., Stubbe, J. Mechanistic studies on bleomycin-mediated DNA damage multiple binding modes can results in double-stranded DNA cleavage Nucleic Acids Research 36, 3781–3790 (2008).

5. Chen, Y., Xu, R., Chen, J., Li, X., He, Q. Cleavage of bleomycin hydrolase by caspase-during apoptosis Oncology Reports 30, 939–944 (2013).

6. Zhang, J. H., Xu, M. DNA fragmentation in apoptosis Cell Research 10, 205–211 (2000).

7. Breaker, R. R. DNA enzyme Nature Biotechnology 15, 427–431 (1997).

8. Borggrafe, J., Victor, J., Rosenbach, H., Viegas, A., Gertzen, C. G. W., Wuebben, C., Kovacs, H., Gopalswamy, M., Riesner, D., Steger, G., Schiemann, O., Gohlke, H., Span, I., Etzkorn, M. Time-resolved structural analysis of an RNA-cleaving DNA catalyst Nature 601, 144–149 (2022).

9. Aranda, J., Terrazas, M., Gomez, H., Villegas, N., Orozco, M. An artificial DNAzyme RNA ligase shows a reaction mechanism resembling that of cellular polymerases Nature Catalysis 2, 544–552 (2019).

10. Lee, Y., Klauser, P. C., Brandsen, B. M., Zhou, C., Li, X., Silvermann, S. K. DNA-catalyzed DNA cleavage by a radical pathway with well-defined products Journal of the American Chemical Society 139, 255–261 (2017).

11. Zhou, C., Avins, J. L., Klauser, P. C., Brandsen, B. M., Lee, Y., Silvermann, S. K. DNA-catalyzed amide hydrolysis Journal of the American Chemical Society 138, 2106–2109 (2016).

12. Camden, A. J., Walsh, S. M., Suk, S. H., Silvermann, S. K. DNA oligonucleotide 3’-phosphorylation by a DNA enzyme Biochemistry 55, 2671-2676 (2016)

13. Chandrasekar, J., Wylder, A., Silvermann, S. K. Phosphoserine lyase deoxyribozymes: DNA-catalyzed formation of dehydroalanine residues in peptides Journal of the American Chemical Society 137, 9575–9578 (2015).

14. Wang, Y., Silvermann, S. K., Directing the outcome of deoxyribozyme selections to favor native 3’-5’ RNA ligation Biochemistry 44, 3017-3023 (2005).

15. Carmi, N., Balkhi, S. R., Breaker R. Cleaving DNA with DNA Proc. Natl. Acad. Sci. USA 95, 2233–2237 (1998).

16. Breaker, R. R., Carmi, N. Characterization of a DNA-cleaving deoxyribozyme Bioorganic & Medicinal Chemistry 9, 2589–2600 (2001).

17. Pogozelski, W. K., Tullius, T. D. Oxidative stress scission of nucleic acids: routes initiated by hydrogen abstraction from the sugar moiety Chemical Reviews 98, 1089–1107 (1998).

18. Gutteridge, J. M. C., Wilkins, S. Copper-dependent hydroxyl radical damage to ascorbic acid Febs Letters 137, 327–330 (1982)

19. Parsekar, S. U., Fernandes, J., Banarjee, A., Chouhan, O. P., Biswas, S., Singh, M., Mishra, D. P., Kumar, M. DNA binding, cleavage and cytotoxicity studies of threemononuclear C(II) chloro-complexes containing N-S donor Schiff base ligands Journal of Biological Inorganic Chemistry 23, 1331–1349 (2018).

20. Yu, W., Wang, S., Cao, D., Rui, H., Liu, C., Sheng, Y., Sun, Y., Zhang, J., Xu, J., Jiang, D. Insight into oxidative DNA-cleaving DNAzyme: multiple cofactors, the catalytic core and a highly efficient variant iScience 23, 101555 (2020).

21. Mattioli, E. J., Bottoni, A., Calvaresi, M. DNAzyme at work: a DFT computational investigation on the mechanism of 9DB1 Journal of Chemical Information and Modeling 59, 1547-1553 (2019).

22. Micura, R., Hobartner, C., Fundamental studies of functional nucleic acids: aptamers, riboswitches, ribozymes and DNAzymes Chemical Society Reviews 49, 7331–7353 (2020).

23. Salvatierra, A. P., Wawrzyniak-Turek, K., Steuerwald, U., Hobartner, C., Pena, V. Crystal structure of a DNA catalyst Nature 529, 231–234 (2016).

24. Joseph, B., Jaumann, E. A., Sikora, A., Barth, K., Prisner, T.F., Cafiso, D. S. In situ observation of conformational dynamics and protein-ligand/substrate interaction in outer membrane proteins with DEER/PELDOR spectroscopy Nature Protocols 14, 2344–2369 (2019).

25. Schiemann, O., Heubach, C. A., Abdullin, D., Ackermann, K., Azarkh, M., Bagryanskaya, E. G., Drescher, M., Endeward, B., Freed, J. H., Galazzo, L., Goldfarb, D., Hett, T., Esteban Hofer, L., Fábregas Ibáñez, L., Hustedt, E. J., Kucher, S., Kuprov, I., Lovett, J. E., Meyer, A., Ruthstein, S., Saxena, S., Stoll, S., Timmel, C. R., Di Valentin, M., Mchaourab, M. S., Prisner, T. F., Bode, B. E., Bordignon, B., Bennati, M., Jeschke, G. Benchmark Test and Guidelines for DEER/ PELDOR Experiments on Nitroxide-Labeled Biomolecules Journal of the American Chemical Society 143, 17875–17890 (2021).

26. Hoebartner, C., Sicoli, G., Wachovius, F., Gophane, D. B., Sigurdsson, S. Th. Synthesis and characterization of RNA containing a rigid and nonperturbing cytidine-derived spin label Journal of Organic Chemistry 77, 7749–7754 (2012).

27. Tkach, I., Pornsuwan, S., Hoebartner, C., Wachowius, F., Sigurdsson, S. Th., Baranova, T. Y., Diederichsen, U., Sicoli, G., Bennati, M. Physical Chemistry Chemical Physics 15, 3433–3437 (2013).

28. Mouesca, J. M., Ahouari, H., Chandra Dantu, S., Sicoli, G. A trade-off for covalent and intercalation binding modes: a case study for Copper (II) ions and singly modified DNA nucleoside Scientific Reports 9, 12602 (2019).

29. Santangelo, M. G., Medina-Molner, A., Schweiger, A., Mitrikas, G., Spingler, B. Structural analysis of Cu(II) ligation to the 5’-GMP nucleotide by pulse EPR spectroscopy Journal of Biological Inorganic Chemistry 12, 767–775 (2007).

30. Godiksen, A., Vennestrom, P. N. R., Rasmussen, S. B., Mossin, S. Identification and quantification of Copper sites in zeolites by electron paramagnetic resonance spectroscopy Topics in Catalysis 60, 13–29 (2017).

31. Prosek, J. Fenton Chemistry in biology and medicine Pure and Applied Chemistry 79, 2325–2338 (2007).

32. Sicoli, G., Mouesca, J.-M., Zeppieri, L., Amara, P., Martin, L., Barra, A.-L., Fontecilla-Camps, J.-C., Gambarelli, S., Nicolet, Y. Fine-tuning of a radical-based reaction by radical S-Adenosyl Methionine (SAM) tryptophan lyase Science 351, 1320-1323 (2016).

33. Bonke, S. A., Risse, T., Schnegg, A., Bruckner, A. In situ electron paramagnetic resonance spectroscopy for catalysis Nature Reviews Methods Primers 1, art. number 33 (2021).

34. Borggrafe, J., Gertzen, C. G. W., Viegas, A., Gohlke, H., Etzkorn, M. The architecture of the 10–23 DNAzyme and its implications for DNA-mediated catalysis FEBS Journal 290, 2011-2021 (2023).

35. Zhou, W., Saran, R., Huang, J.-P. J., Ding, J., Liu, J. An exceptionally selective DNA cooperatively binding two Ca 2+ ions ChemBioChem 18, 518–522 (2017).

36. Huang, P.-J. J., Vazin, M., Liu, J. In vitro selection of a DNAzyme cooperatively binding two lanthanides ions for RNA cleavage Biochemistry 55, 2518–2525 (2016).

37. Cao, Y., Zhang, H., Le, X. C. Split locations and secondary structures of a DNAzyme critical to binding-assembled multicomponent nucleic acid enzymes for protein detection Analytical Chemistry 93, 15712–15719 (2021).

38. Zhou, W., Zhang, Y., Huang, P.-J. J., Ding, J., Liu, J. A DNAzyme requiring two different metal ions at two distinct sites Nucleic Acids Research 44, 354–363 (2016).

39. Zhank, Y., Ji, Z., Wang, X., Cao, Y., Pan, H. Single-molecule study of DNAzyme reveals its intrinsic conformational dynamics International Journal of Molecular Sciences 24, art. n. 1212 (2023).

40. Rosenbach, H., Victor, J., Etzkorn, M., Steger, G., Riesner, D., Span, I. Molecular features and metal ions that influence 10–23 DNAzyme activity Molecules 25, 3100 (2020).

## References

41. Lu, X.-J., Olson, W. K. 3DNA: a software package for the analysis, rebuilding and visualization of three-dimensional nucleic acid structures Nuclei Acids Research 31, 5108-5121 (2003).

42. Ulyanov, N. B., Bauer, W. R., James, T. L. High-resolution NMR structure of an AT-rich DNA sequence Journal of Biomolecular NMR 22, 265–280 (2002).

43. Iyani, I., Dans, P. S., Noy, A., Perez, A., Faustino, I., Hospital, A., Walther, J., Andrio, P., Goni, R., Balaceanu, A., Portella, G., Battistini, F., Gelpi, J. L., Gonzales, C., Vendruscolo, M., Laughton, C. A., Harris, S. A., Case, D. A., Orozco, M. Parmbsc1: a refined force field for DNA simulations Nature Methods 13, 55–58 (2016).

44. Abraham, M. J., Murtola, T., Schulz, R., Pall, S., Smith, J. C., Hess, B., Lindahl, E. GROMACS: high performance molecular simulations through multi-level parallelism from laptops to supercomputers SoftwareX 1-2, 19–25 (2015).

45. Jorgensen, W., Chandrasekhar, J., Madura, J. D., Impey, R. W., Klein, M. L. Comparison of simple potential functions for simulating liquid water Journal of Chemical Physics 79, 926–935 (1983).

46. Cele, F. N., Kumalo, H., Soliman, M. E. S. Mechanism of inhibition of Hsp90 dimerization by inhibitor coumermycin A1 (C-A1) revealed by dynamics simulations and thermodynamic calculations Cell Biochemistry and Biophysics 74, 353–363 (2016).

47. Bussi, G., Donadio, D., Parrinello, M. Canonical sampling through velocity rescaling Journal of Chemical Physics 126, 014101 (2007).

48. Parrinello, M., Rahman, A. Polymorphic transitions in single crystals: a new molecular dynamics method Journal of Applied Physics 52, 7182–7190 (1981).

49. Daura, X, Gademan, K., Jaun, B., Seebach, D., van Gunsteren, W. F., Mark, A. E. Peptide folding: when simulation meets experiment Angewandte Chemie International Edition 38, 236–240 (1999).

50. PyMOL (RRID:SCR_000305), Molecular Graphics System, Version 2.0 Schrödinger, LLC.

51. Sacco, A, Heil, S, Self-Diffusion coefficients D of various liquids at 25°C. Courtesy (Éd.), Almanach 80 (2007).

52. Collectif. Diffusion Studied Using NMR Spectroscopy. E. Science, Éd (2016).

53. Haruhisa, K., Takeshi, S, Mami, N., Kayori, S., Shinichi, K., Assessment of diffusion coefficients of general solvents by PFG-NMR: Investigation of the sources error, Journal of magnetic Resonance 180, 266–273 (2006).

54. Holz, M., Weingartner, H., Calibration in Accurate Spin-Echo Self-Diffusion Measurements Using 1H and Less-Common Nuclei Journal of Magnetic Resonance 92, 115–125 (1991).

55. Holz, M, Heil, S, Sacco, A, Temperature dependent self-diffusion coefficients of water and six selected molecular liquids for calibration in accurate 1H NMR PFG measurements. Phys. Chem. Chem. Phys 2, 4740–4742. (2000).

56. Kerssebaum, R, Salnikov, G,. Dosy and diffusion by NMR. BRUKER. A tutorial for Topspin 2.0.0 Version 2.0.0 (2002-2006).

57. Guo, B. Mass spectrometry in DNA analysis Analytical Chemistry 71, 333–337 (1999).

58. Capobianco, A., Caruso, T., D’Ursi, A. M., Fusco, S., Masi, A., Scrima, M., Chatgilialoglu, C., Peluso, A. Delocalized hole domains in guanine-rich DNA oligonucleotides The Journal of Physical Chemistry B 119, 5462–5466 (2015).

59. Taylor, A. I., Pinheiro, V. B., Smola, M. J., Morgunov, A. S., Chew, A. S., Cozens, C., Weeks, K. M., Herdewiin, P., Holliger, P. Catalysts from synthetic genetic polymers Nature 18, 427–430 (2015).

