## Supplementary Information for "Cleaving DNA with DNA: cooperative tuning of structure and reactivity driven by copper ions"

### Figure S1

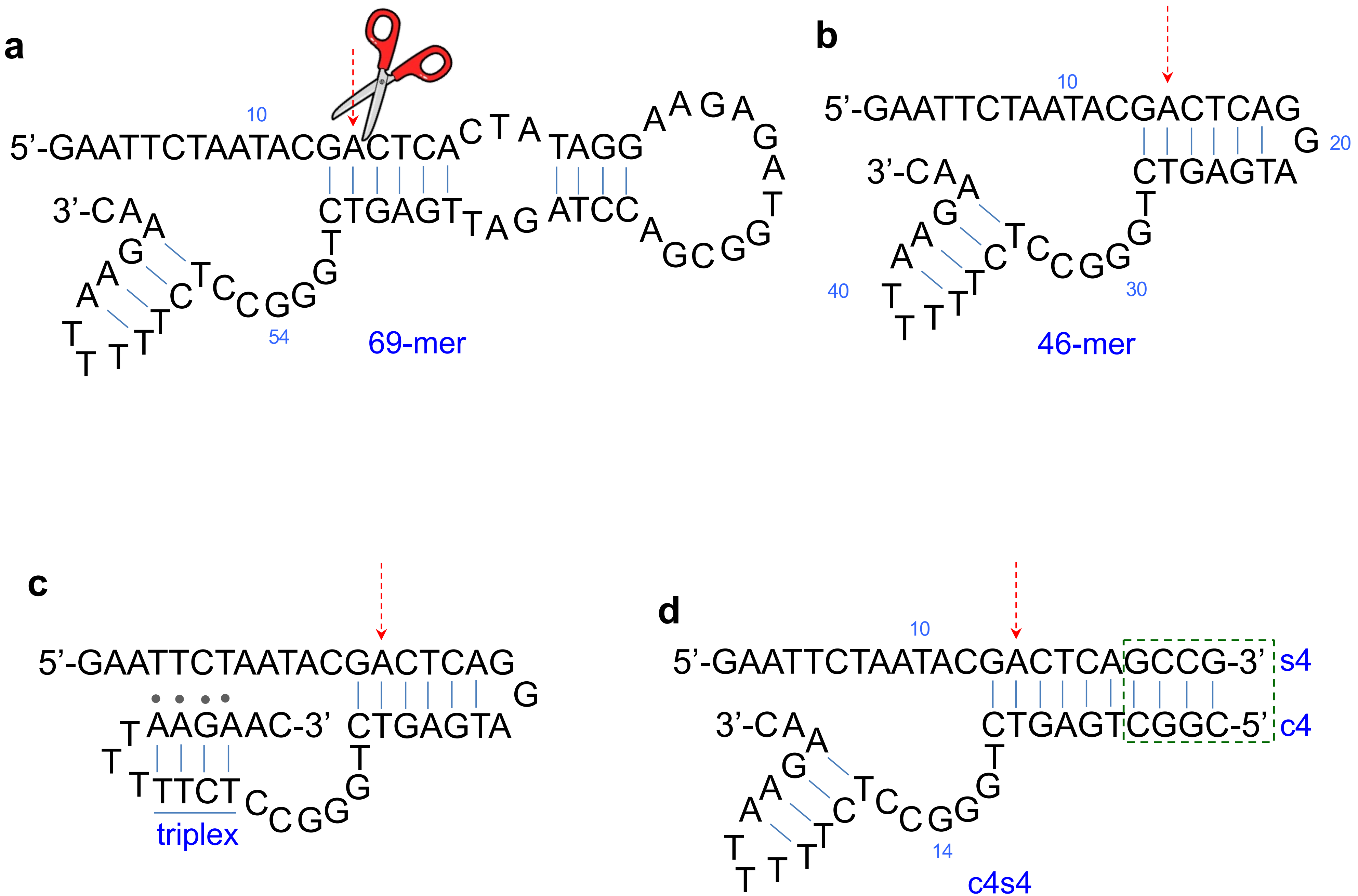
a
10
5’-GAATTCTAATACGACTCA
CTA
TAGG
AGA
CTGAGT
TAG
ATCC
3’-CA
54
69-mer
b
10
5’-GAATTCTAATACGACTCAG
G
20
CTGAGTA
3’-CA
40
30
46-mer
c
5’-GAATTCTAATACGACTCAG
G
AAGAAC-3’
CTGAGTA
TTCT
triplex
d
10
5’-GAATTCTAATACGACTCAGCCG-3’
s4
CTGAGTCGGC-5’
c4
3’-CA
14
c4s4

**Figure S2**

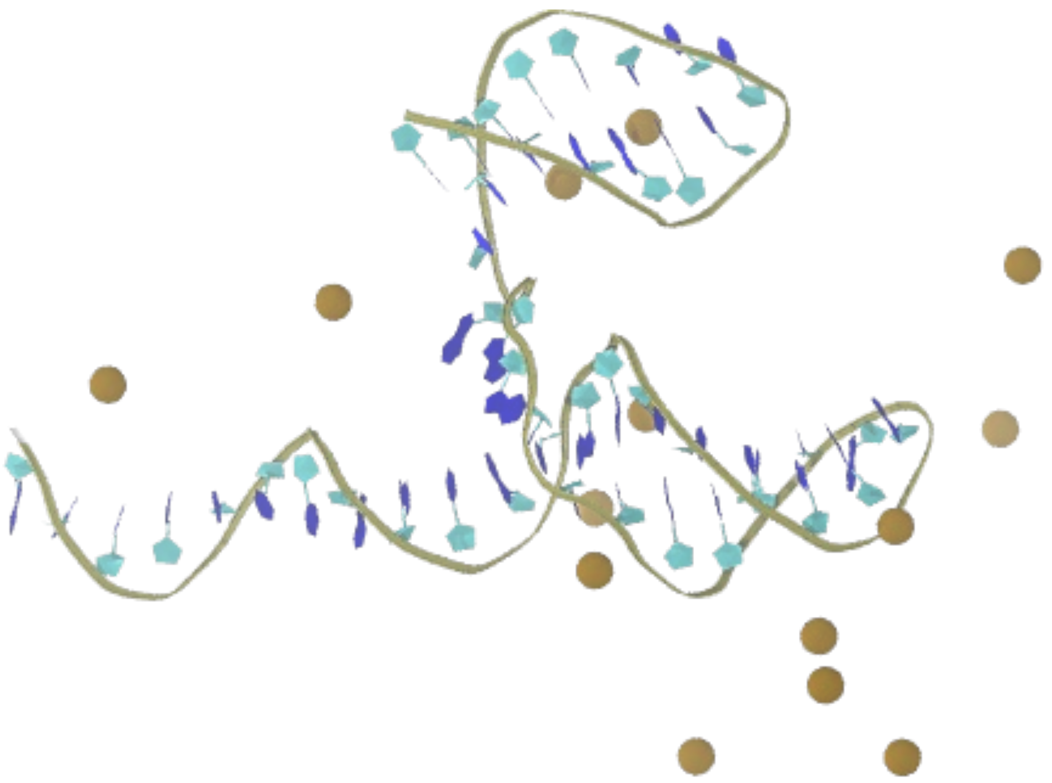

### Figure S3

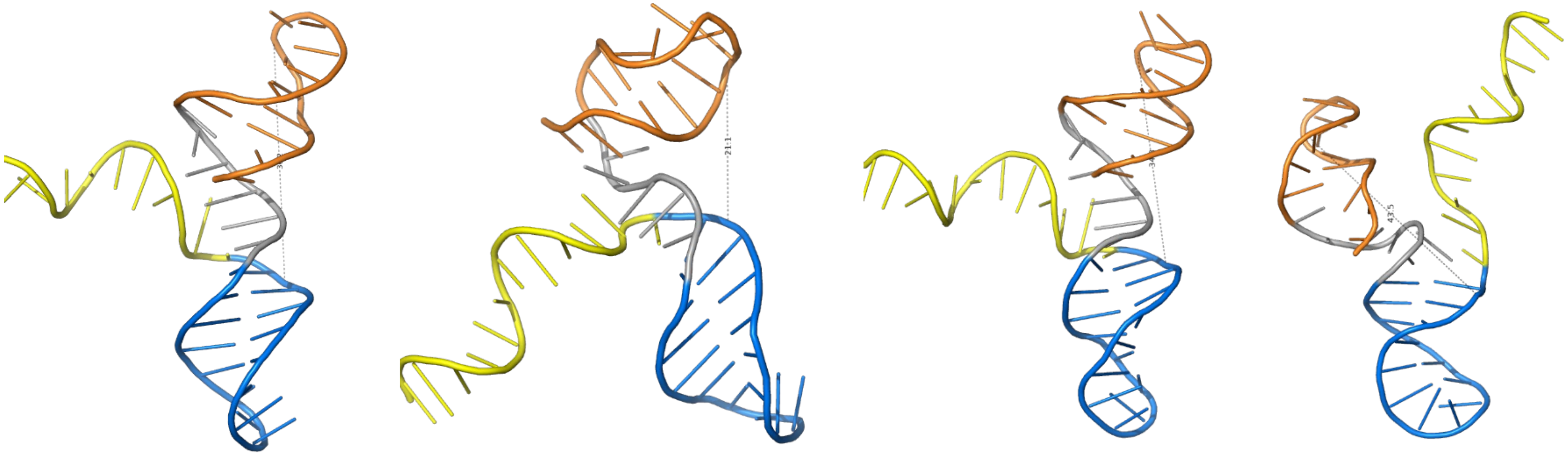

99.4% 0.003% 0.002% 0.001%

### Figure S4

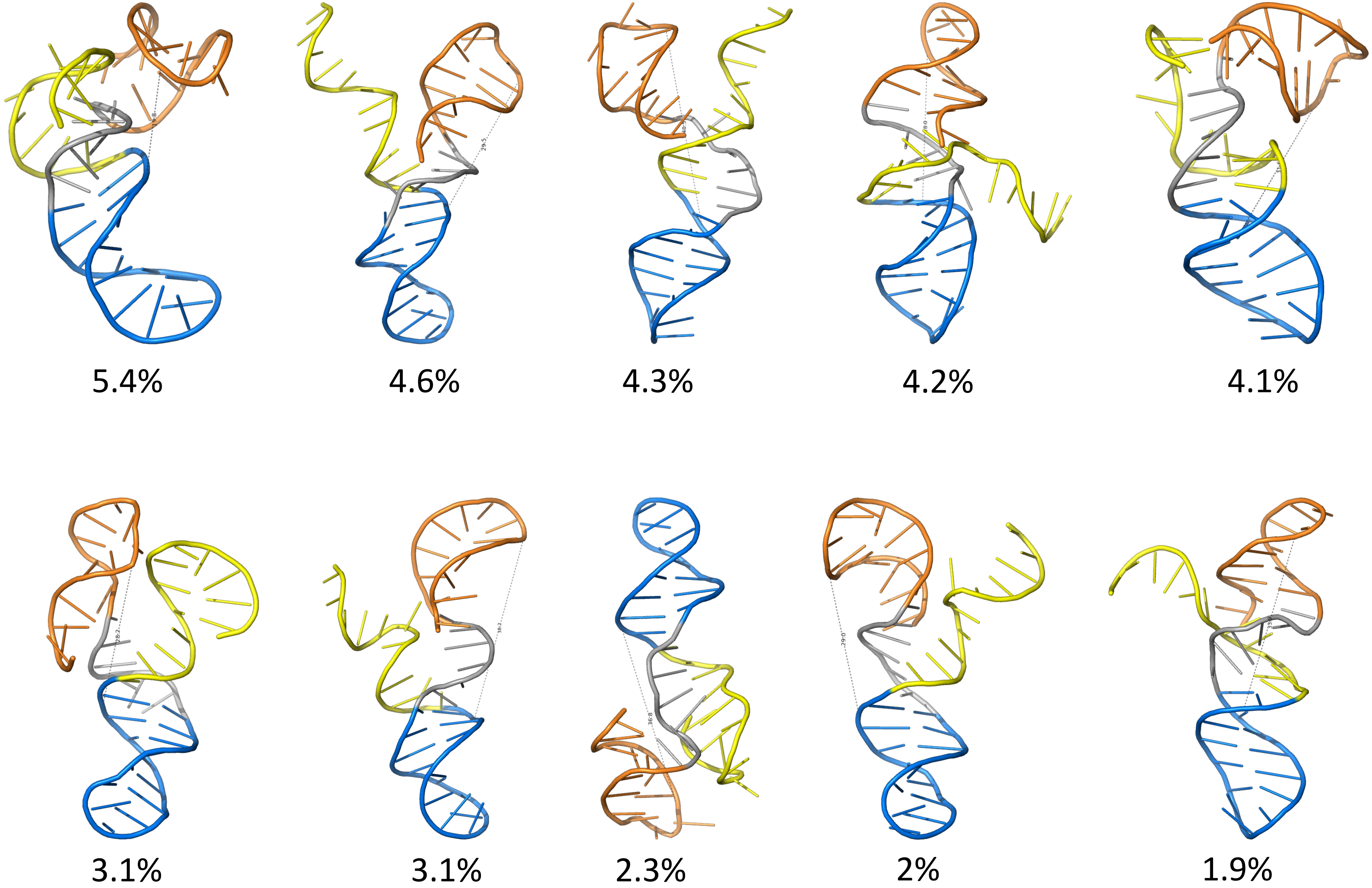

#### Figure S5

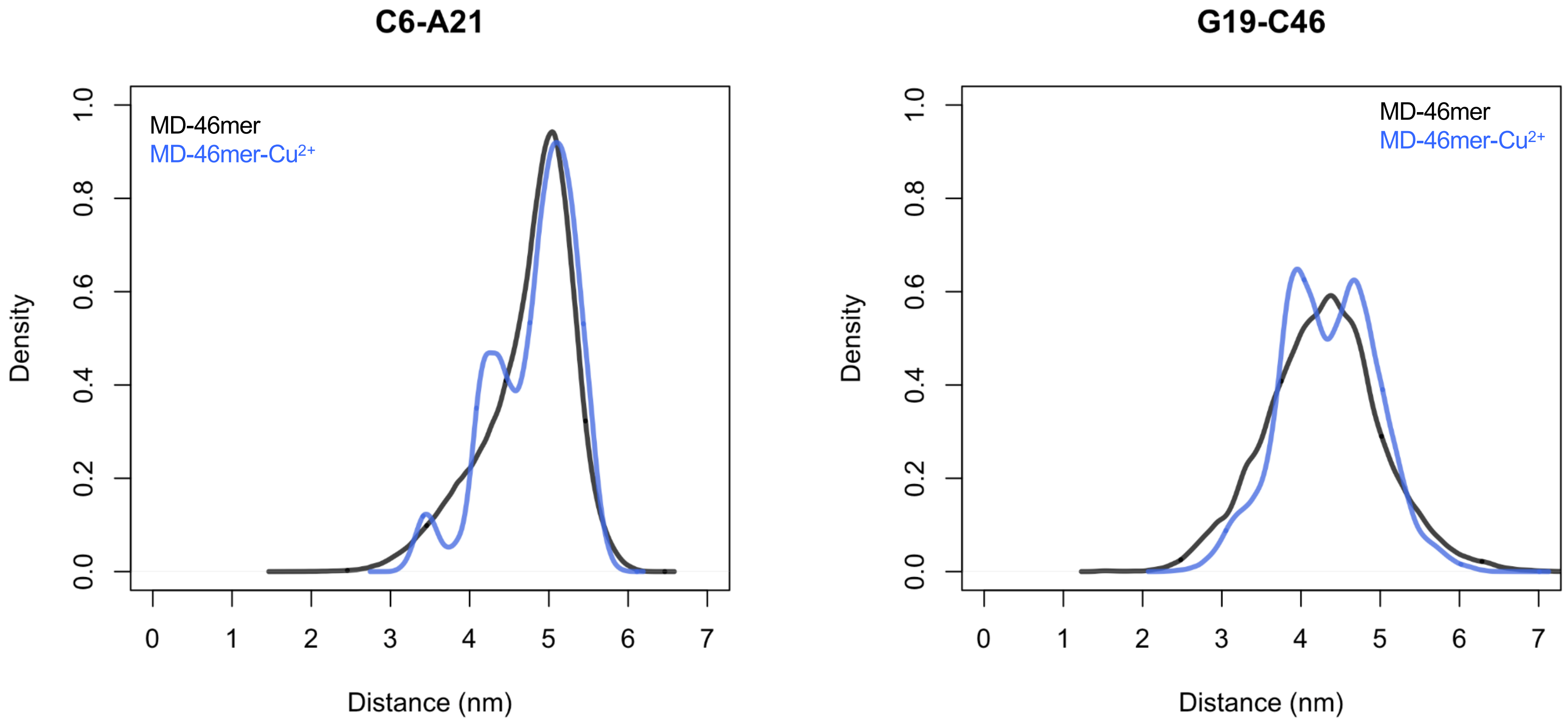
C6-A21
G19-C46
MD-46mer
MD-46mer-Cu2+
Density
Distance (nm)
0.0
0.2
0.4
0.6
0.8
1.0
0
1
2
3
4
5
6
7

### Figure S6-a

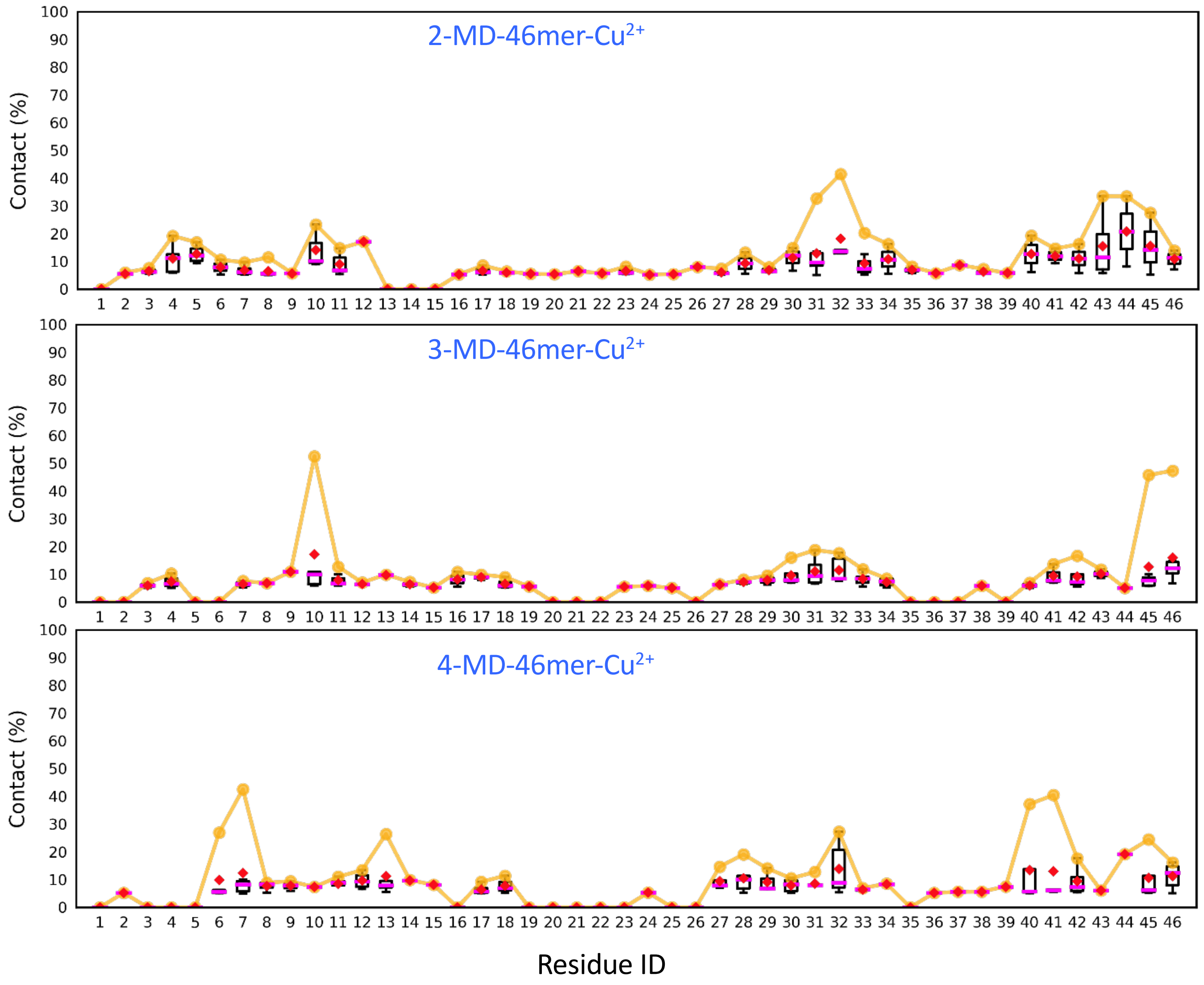

2-MD-46mer-Cu2+
3-MD-46mer-Cu2+
4-MD-46mer-Cu2+
Contact (%)
Residue ID
0
10
20
30
40
50
60
70
80
90
100
1 2 3 4 5 6 7 8 9 10 11 12 13 14 15 16 17 18 19 20 21 22 23 24 25 26 27 28 29 30 31 32 33 34 35 36 37 38 39 40 41 42 43 44 45 46

### Figure S6-b

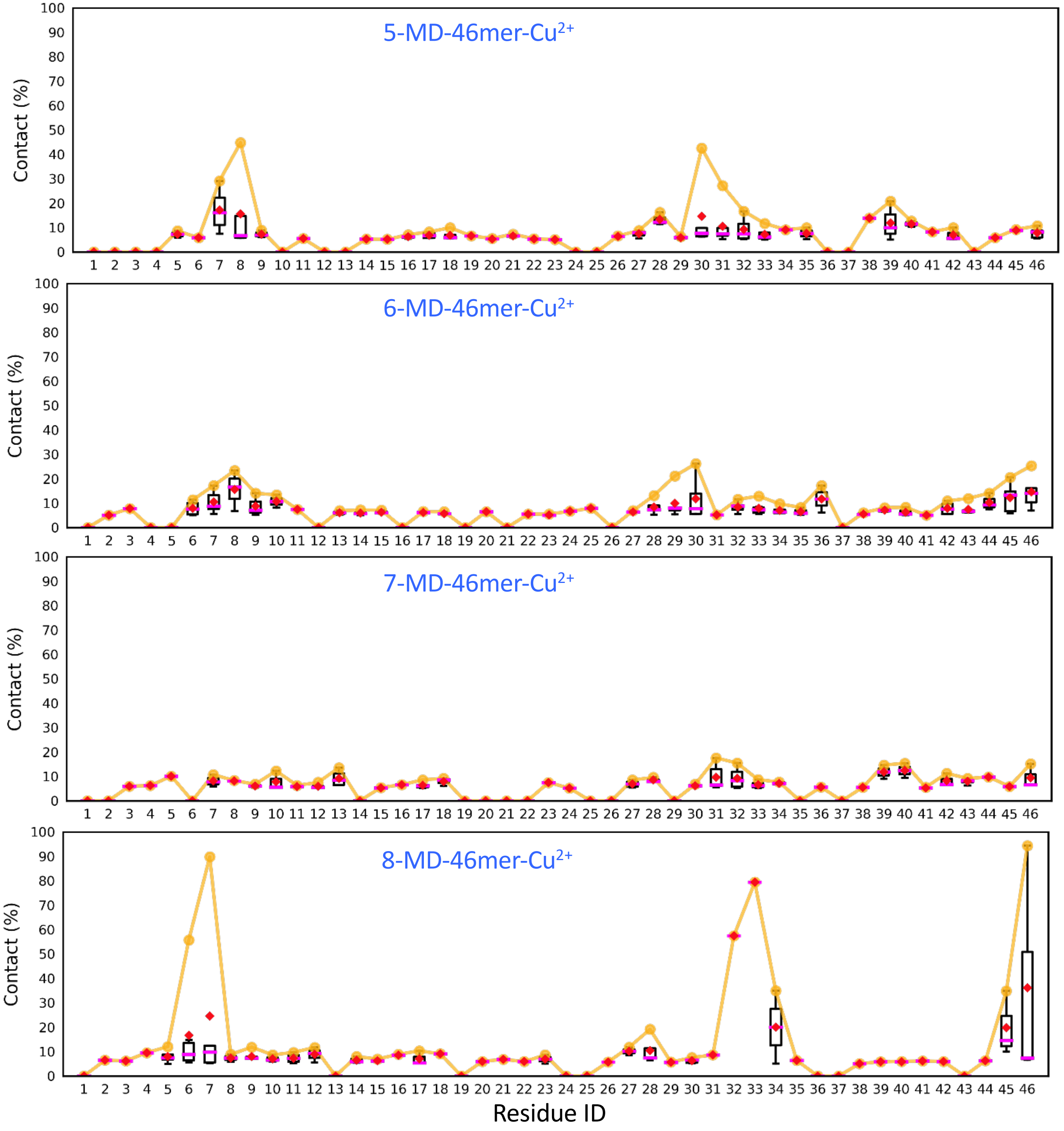

5-MD-46mer-Cu2+
6-MD-46mer-Cu2+
7-MD-46mer-Cu2+
8-MD-46mer-Cu2+
Contact (%)
Residue ID

### Figure S6-c

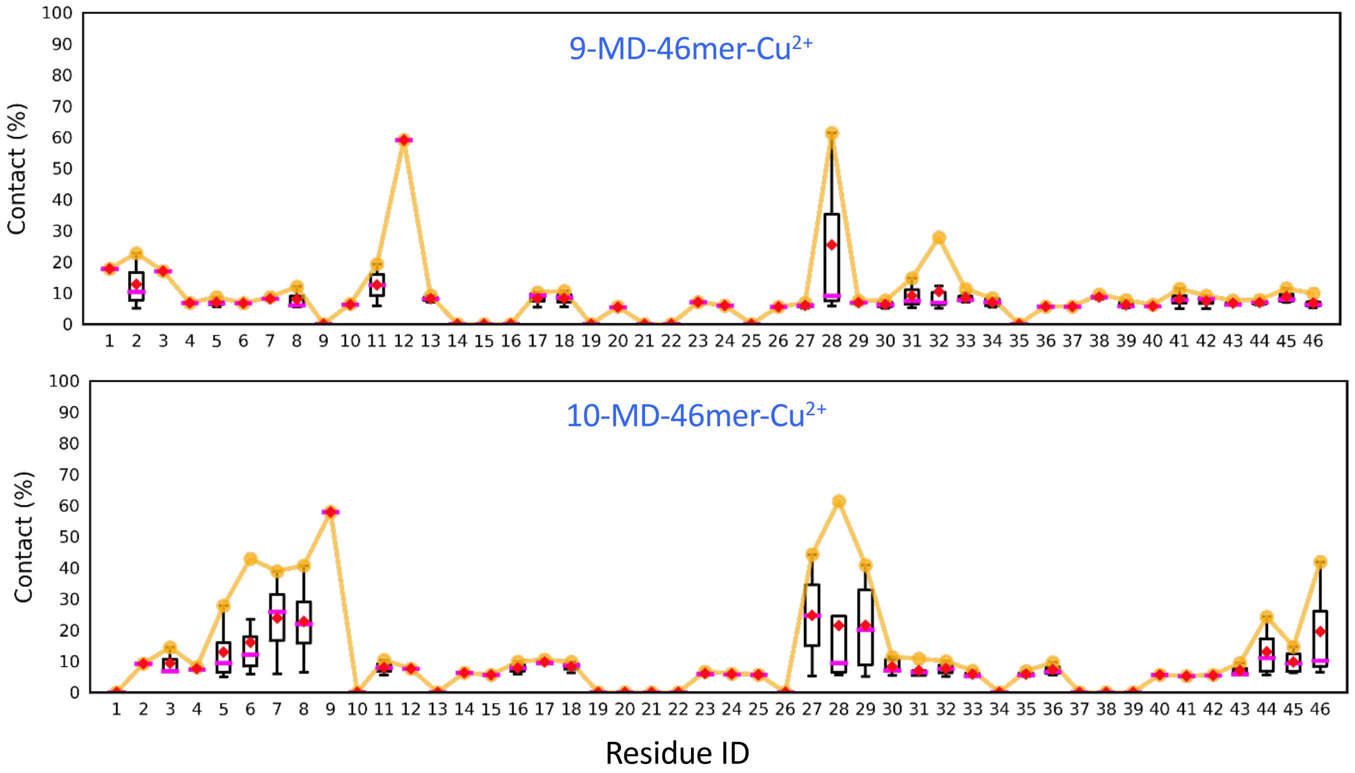

### Figure S7a

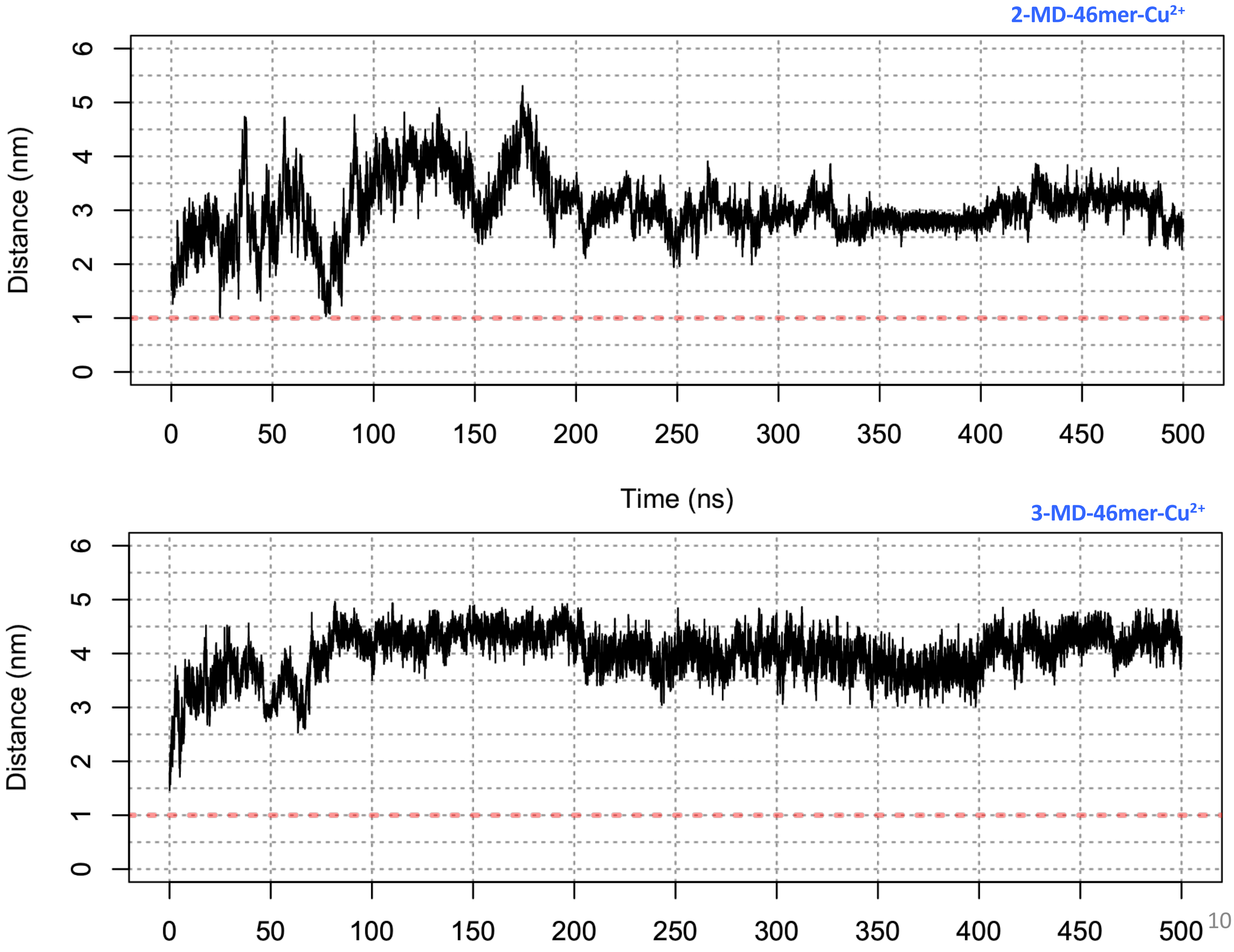

2-MD-46mer-Cu2+
Distance (nm)
0
1
2
3
4
5
6
0
50
100
150
200
250
300
350
400
450
500
Time (ns)
3-MD-46mer-Cu2+
Distance (nm)
0
1
2
3
4
5
6
0
50
100
150
200
250
300
350
400
450
500

### Figure S7b

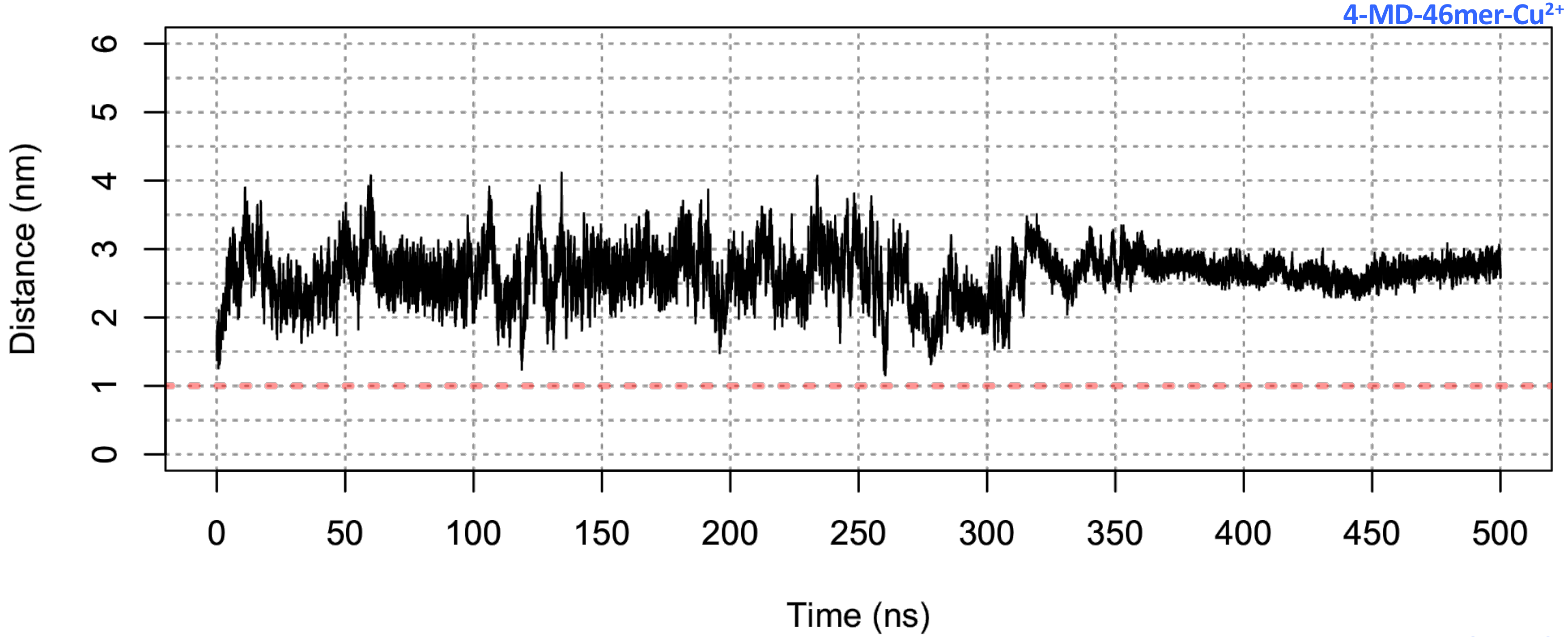
4-MD-46mer-Cu2+
Distance (nm)
0
1
2
3
4
5
6
0
50
100
150
200
250
300
350
400
450
500
Time (ns)

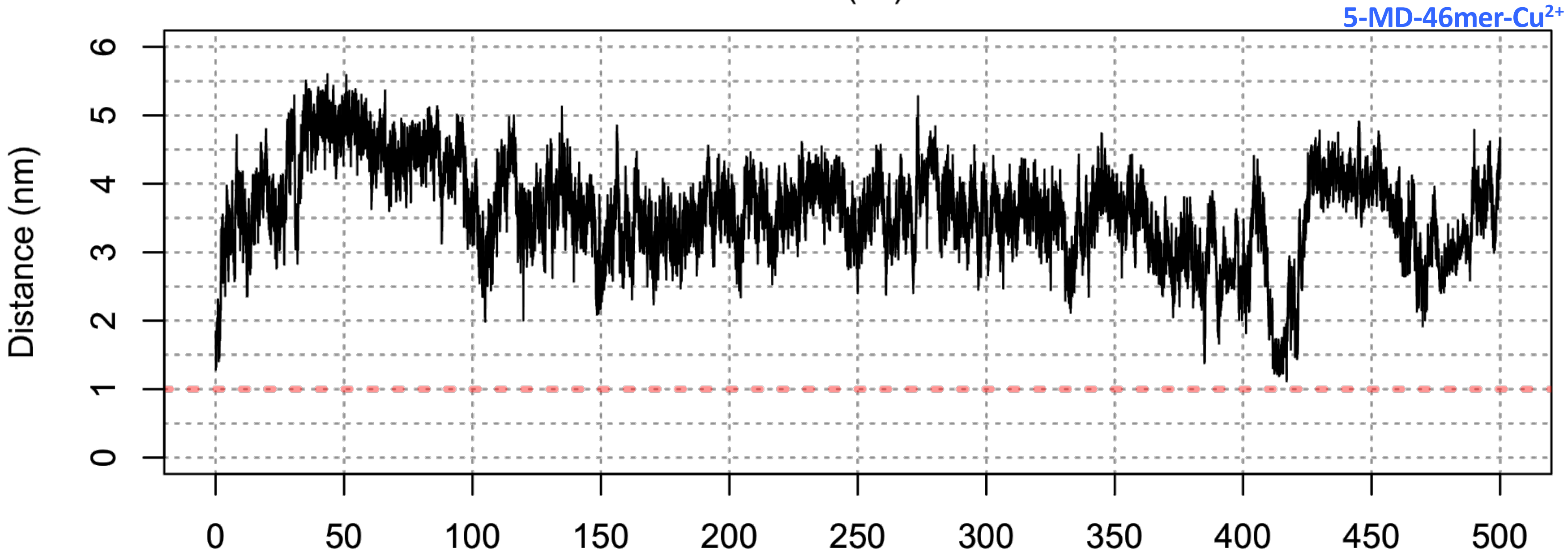
5-MD-46mer-Cu2+
Distance (nm)
0
1
2
3
4
5
6
0
50
100
150
200
250
300
350
400
450
500

### Figure S7c

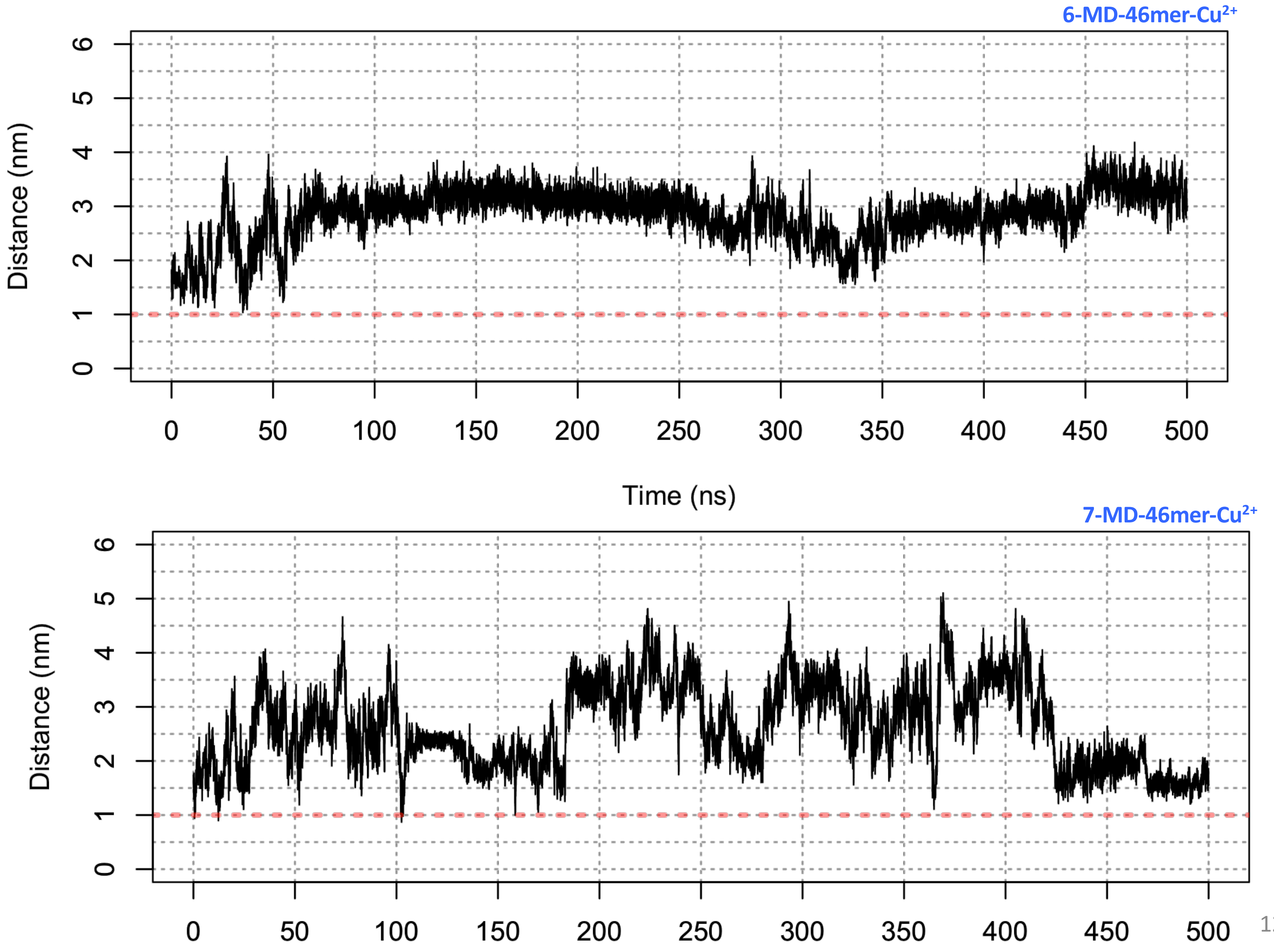
6-MD-46mer-Cu2+
Distance (nm)
0
1
2
3
4
5
6
0
50
100
150
200
250
300
350
400
450
500
Time (ns)
7-MD-46mer-Cu2+
Distance (nm)
0
1
2
3
4
5
6
0
50
100
150
200
250
300
350
400
450
500

### Figure S7d

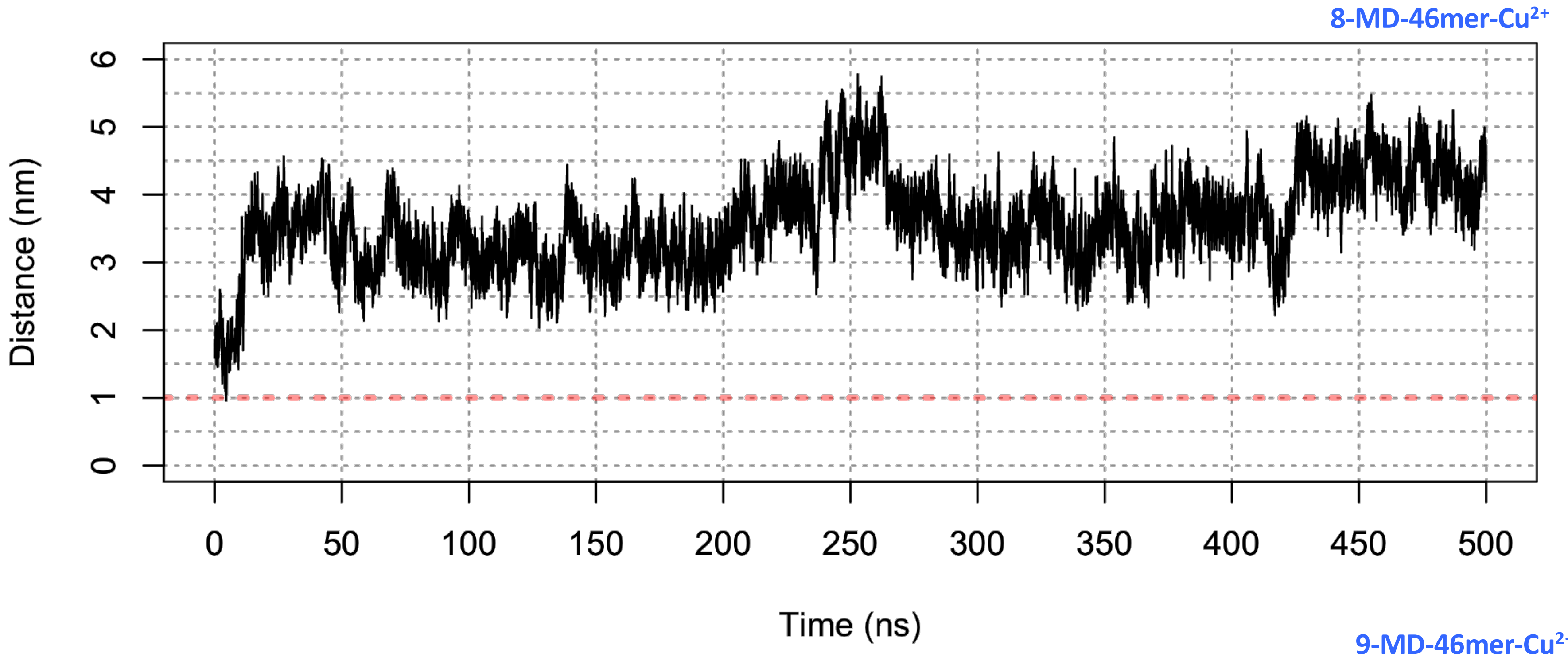
8-MD-46mer-Cu2+
Distance (nm)
0
1
2
3
4
5
6
0
50
100
150
200
250
300
350
400
450
500
Time (ns)

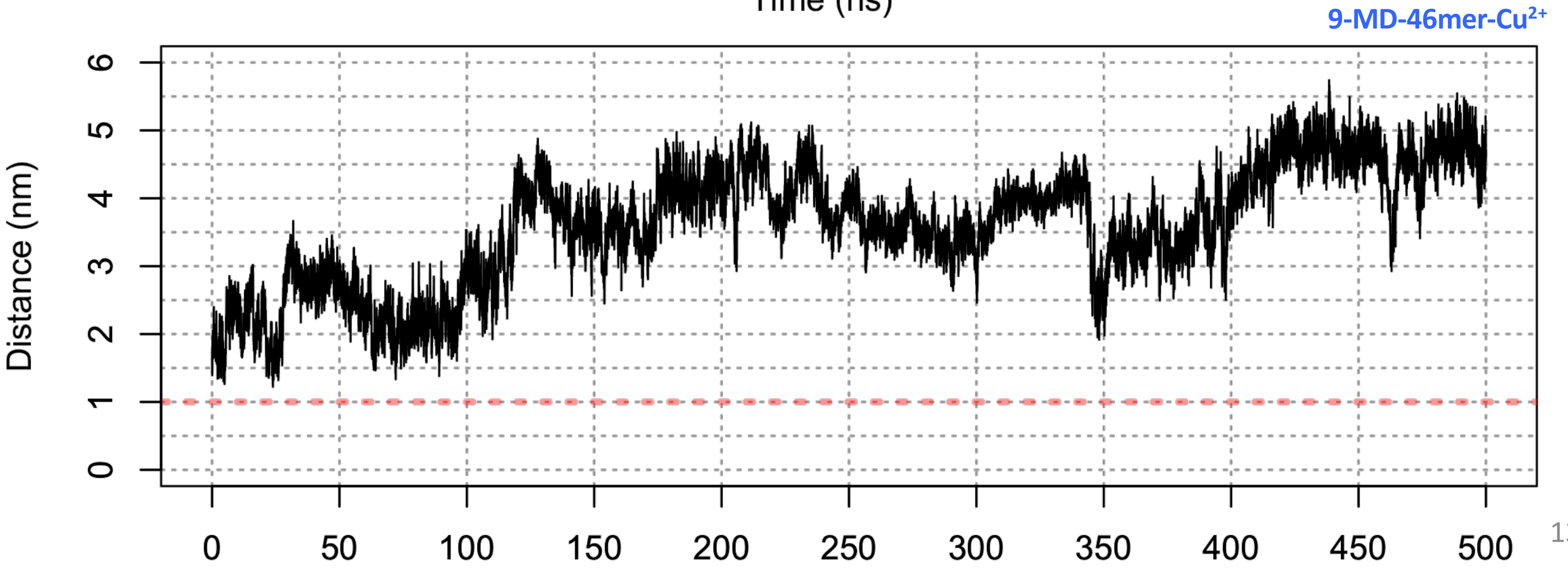
9-MD-46mer-Cu2+
Distance (nm)
0
1
2
3
4
5
6
0
50
100
150
200
250
300
350
400
450
500

### Figure S7e

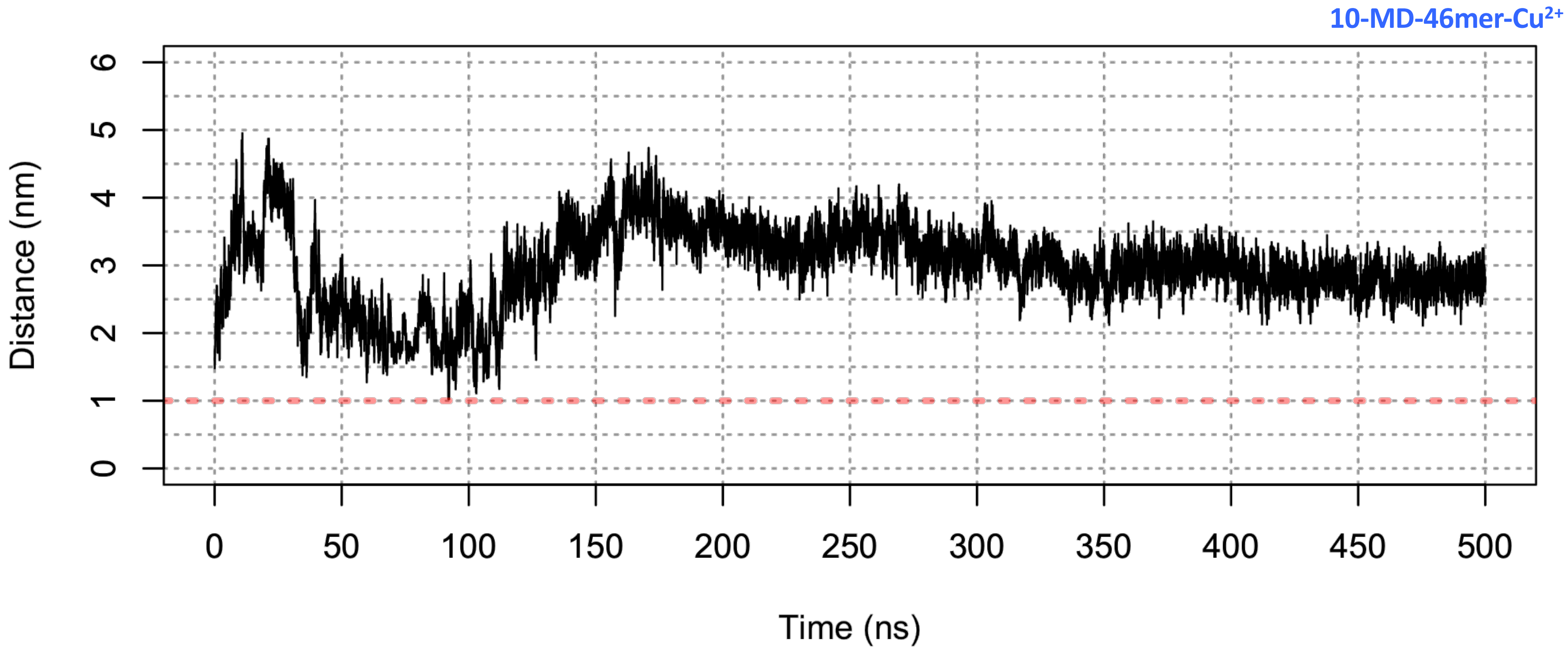

10-MD-46mer-Cu2+
Distance (nm)
0
1
2
3
4
5
6
0
50
100
150
200
250
300
350
400
450
500
Time (ns)

### Figure S8

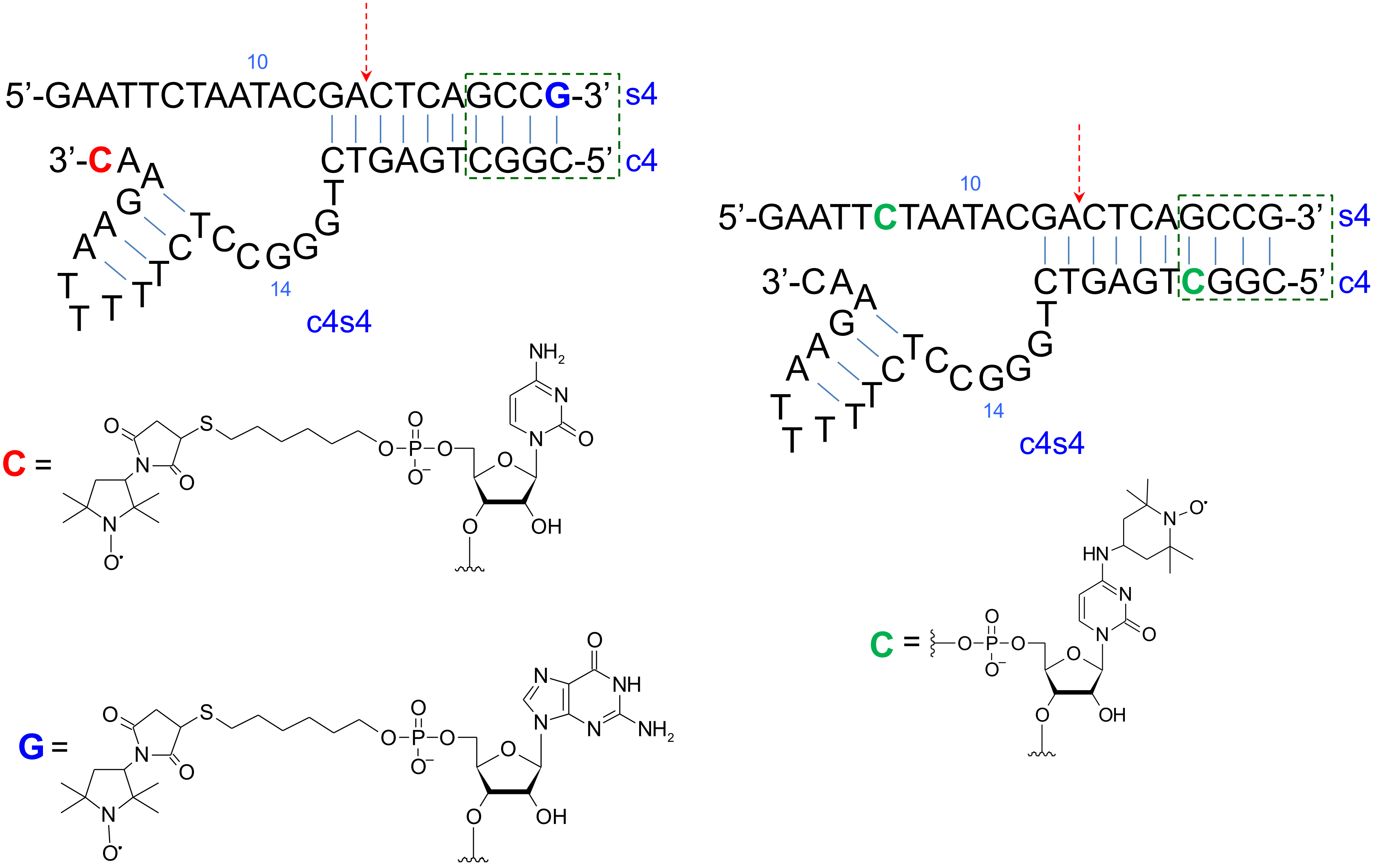

10
5’-GAATTCTAATACGACTCAGCCG-3’ s4
3’-CAAGAATTTTTCTCCGGGTCTGAGTCGGC-5’ c4
14
c4s4
C =
G =
10
5’-GAATTCTAATACGACTCAGCCG-3’ s4
3’-CAAGAATTTTTCTCCGGGTCTGAGTCGGC-5’ c4
14
c4s4
C =

### Figure S9

c4s4_G14(c4)_$^{15}$N

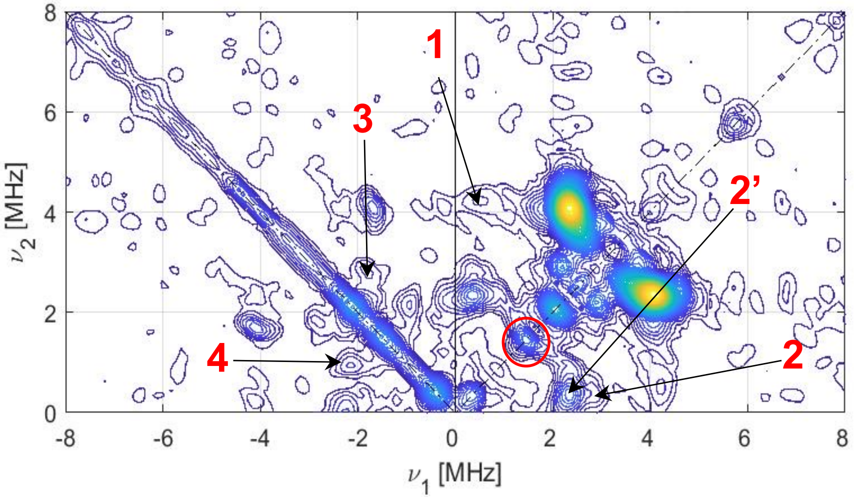


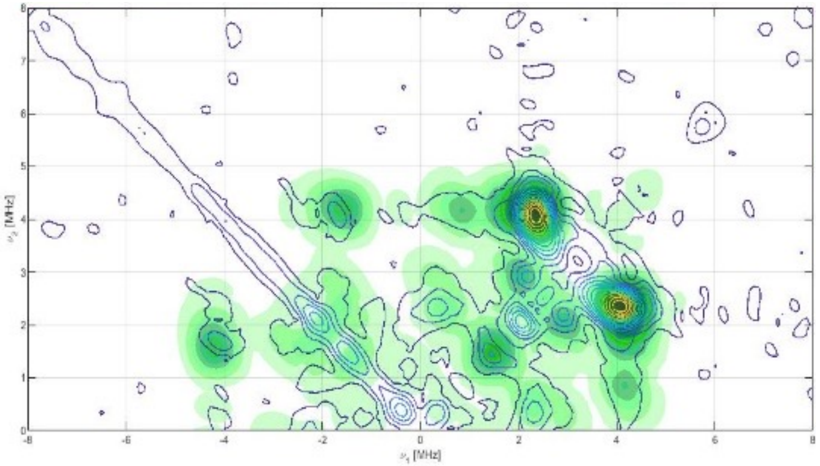

### Table S10

| Samples | $\Delta v\ of (+/+) region\ in\ MHz$ | | | $\Delta v\ of (-/+) region\ in\ MHz$ | | |
|---|---|---|---|---|---|---|
| | 1 | 2 | 2’ | 3 | 4 | 4’ |
| **c4s4** | 3.37 | 2.44 | — | 1.08 | 1.08 | — |
| **C4S4-c4-GUA14-$^{15}$N** | 3.66 | 2.44 | 1.89 | 1.08 | 1.20 | — |
| **C4S4-s4-GUA13-$^{15}$N** | 3.42 | 2.44 | 1.85 | 0.82 | 1.08 | 1.67 |

* $\Delta v = v_2 - v_1$

By comparing the +/+ quadrant of the unlabeled sample to the $^{15}N$ labeled sample at its guanosine number 14 of c4 sequence and the labeled sample at guanosine number 13 of s4 an additional cross peak “ 2’ ” of $\Delta v = 2.7\ MHz$ is observed, as shown in fig.3 c & e. This new cross peak appears beside peak number “ 2 ” in the unlabeled sample (of $\Delta v = 3.45\ MHz$ which is originated by $^{14}N$ double quanta transition). The cross peak “ 2’ ” is characterized with $^{15}N$ Larmor frequency (1.506 MHz) which confirm the presence of $Cu^{2+}$ coordination to $^{15}N$ of the bases G13 and G14 of s4 and c4 respectively. In addition, the comparison between the -/+ quadrant of the unlabeled sample and the labeled one at G14 fig.3 a & c respectively, shows same $\Delta v$ of all the peaks except the peak number “ 4 ” ($\Delta v = 1.48 MHz$) for the unlabeled less than the G14 labeled ($\Delta v = 1.70 MHz$). This broadening can be assigned for coupling to $^{15}N$ (replacing one of the 14N’s).

### Figure S11

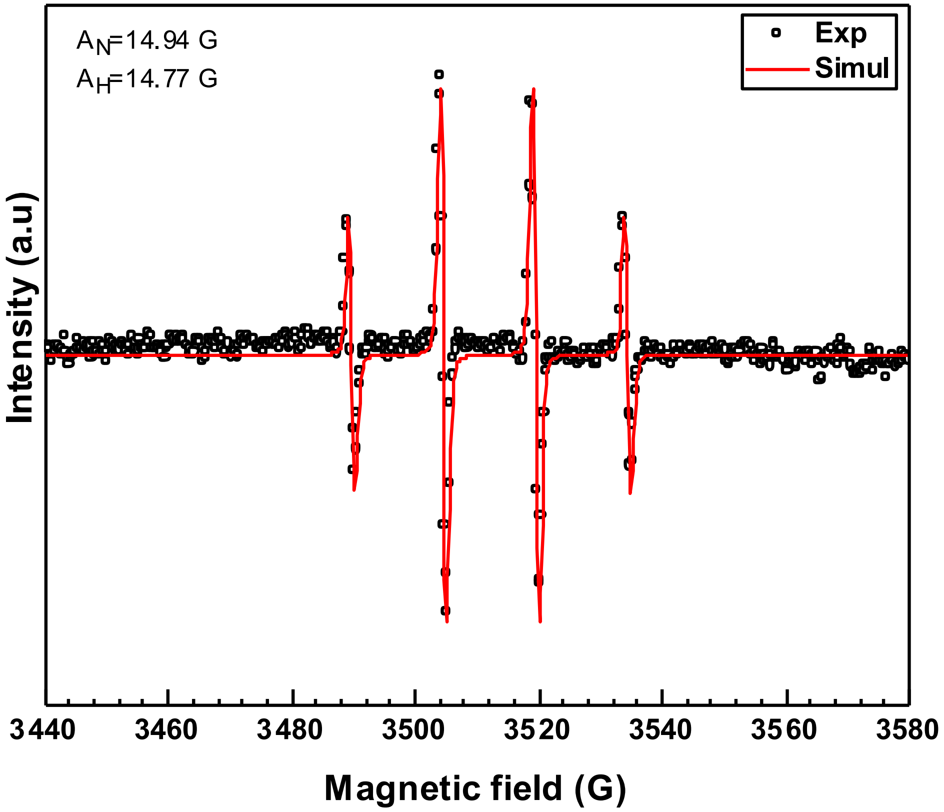

$A_N$=14.94 G
$A_H$=14.77 G
Exp
Simul
Intensity (a.u)
3440
3460
3480
3500
3520
3540
3560
3580
Magnetic field (G)

**Figure S12**

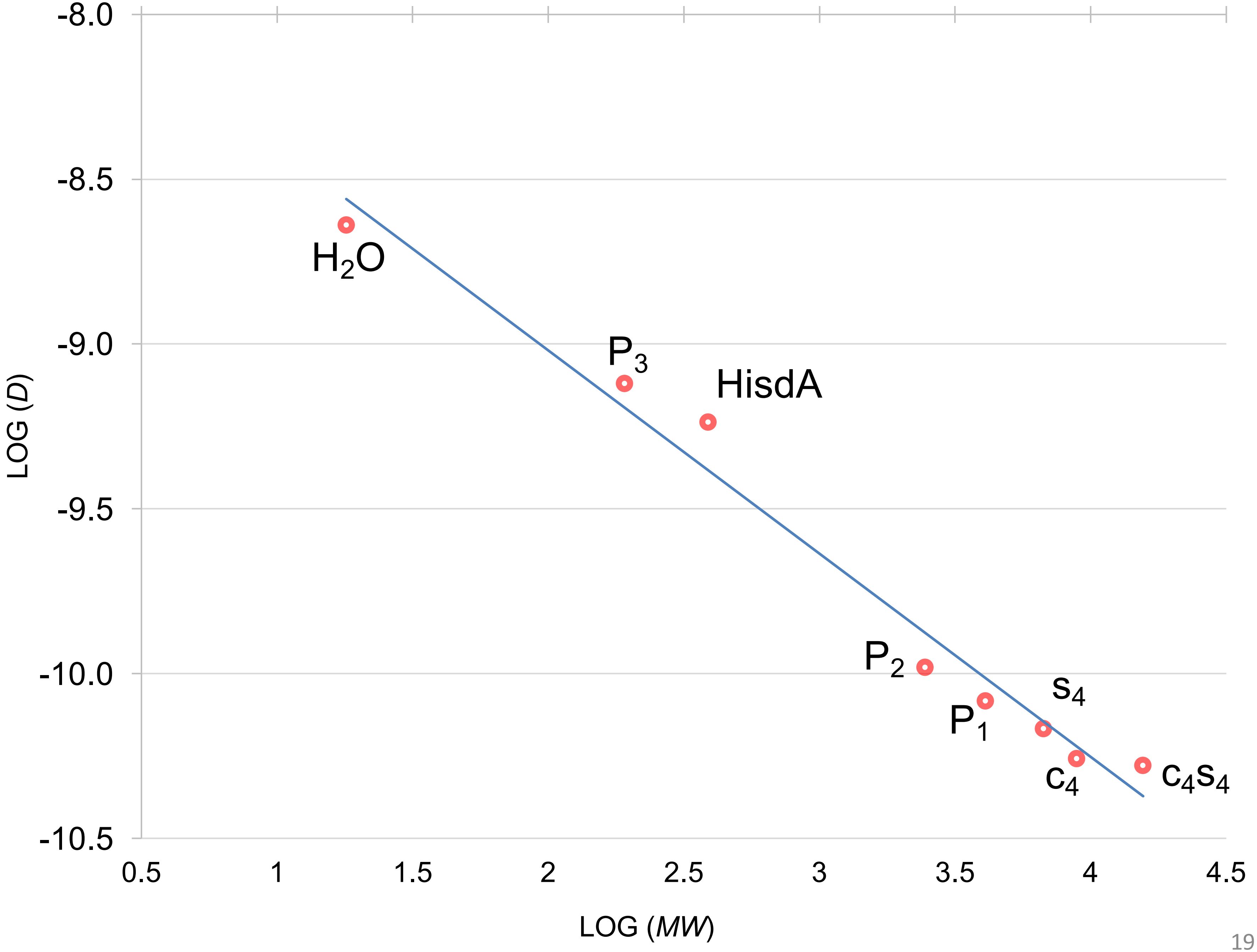
-8.0
-8.5
-9.0
-9.5
-10.0
-10.5
LOG (D)
H2O
P3
HisdA
P2
P1
s4
c4
c4s4
0.5
1
1.5
2
2.5
3
3.5
4
4.5
LOG (MW)

### Table S13

Table S12.

| | $D$(μm$^2$/s) | $MW$(g/mol) | Log($D$) (a.u.) | Log($MW$) (a.u.) |
|---|---|---|---|---|
| **$H_2O$** | 2.3 x $10^{-9}$ | 18 | – 8.64 | 1.26 |
| **$c_4$** | 55.1 x $10^{-12}$ | 8897 | – 10.3 | 3.95 |
| **$s_4$** | 68.1 x $10^{-12}$ | 6704 | – 10.2 | 3.83 |
| **$c_4s_4$** | 52.6 x $10^{-12}$ | 15591 | – 10.3 | 4.19 |
| **His-dA** | 579.3 x $10^{-12}$ | 388 | – 9.24 | 2.56 |
| **$P_1$** | 82.6 x $10^{-12}$ | 4096 | – 10.1 | 3.61 |
| **$P_2$** | 104.5 x $10^{-12}$ | 2451 | – 9.98 | 3.39 |
| **$P_3$** | 757.6 x $10^{-12}$ | 191 | – 9.12 | 2.28 |

### Figure S14

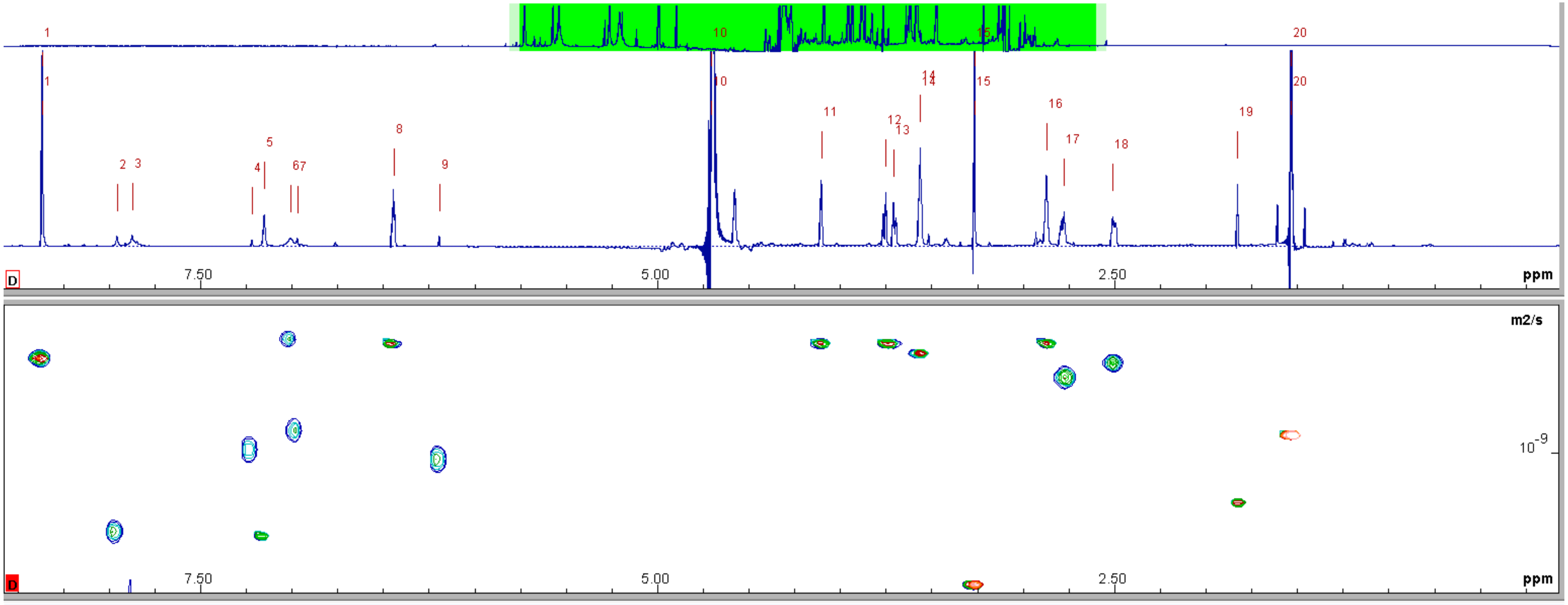


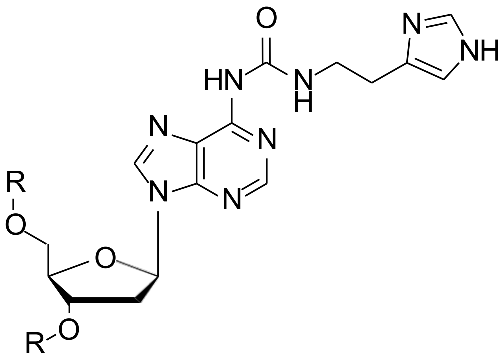

R = H

### Figure S15

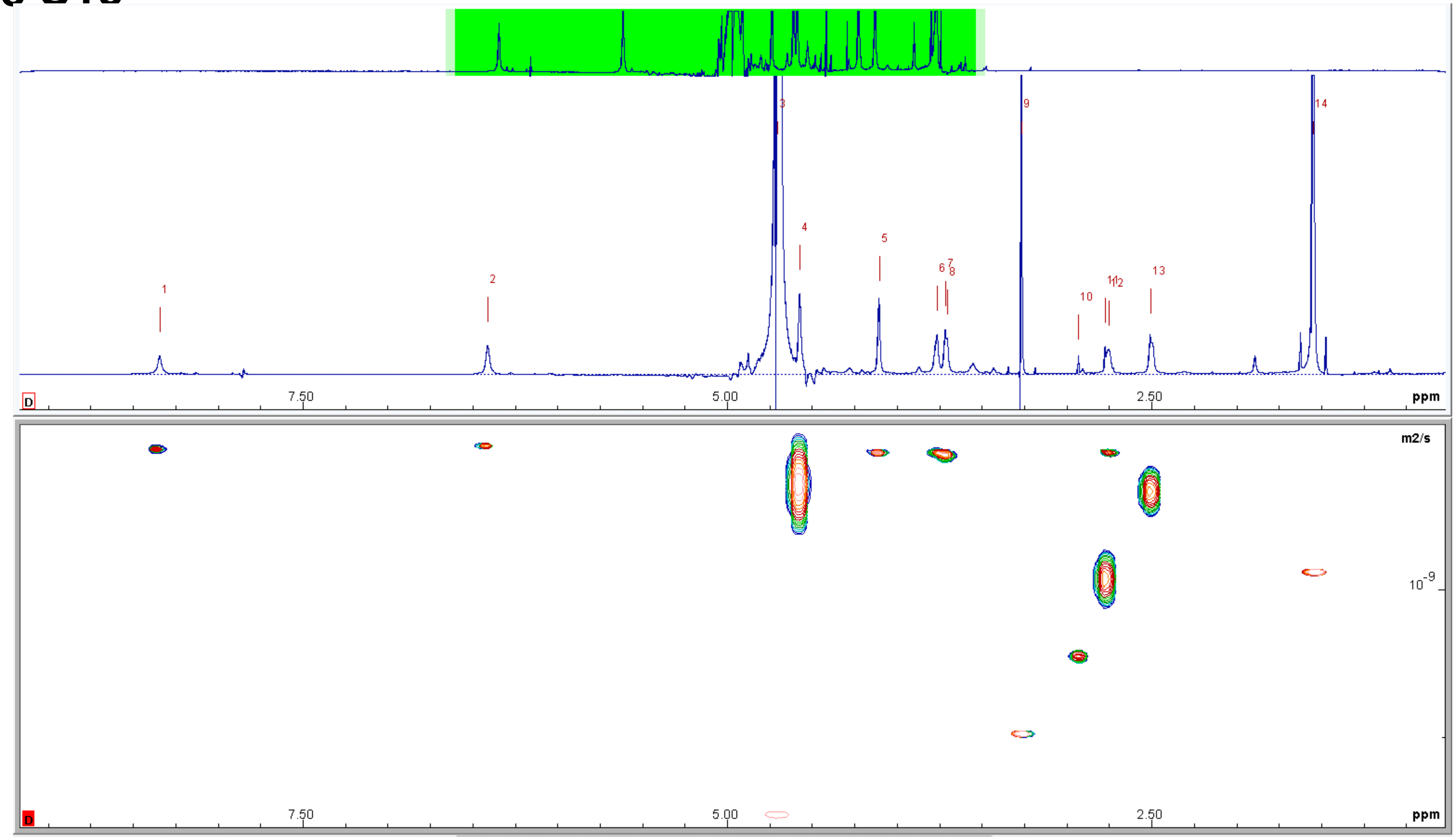

1
2
3
4
5
6
7
8
9
10
11
12
13
14
7.50
5.00
2.50
ppm
m2/s
$10^{-9}$


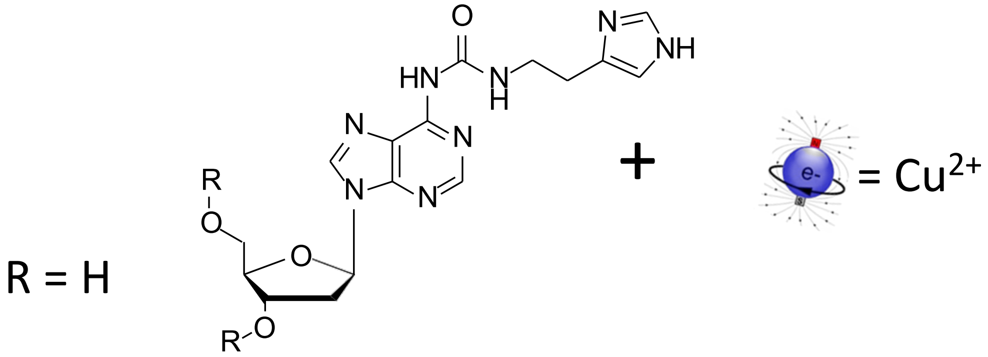

R = H
O
HN
N
H
N
NH
N
N
N
N
R
O
O
R
O
+
e-
= $Cu^{2+}$

### Figure S16

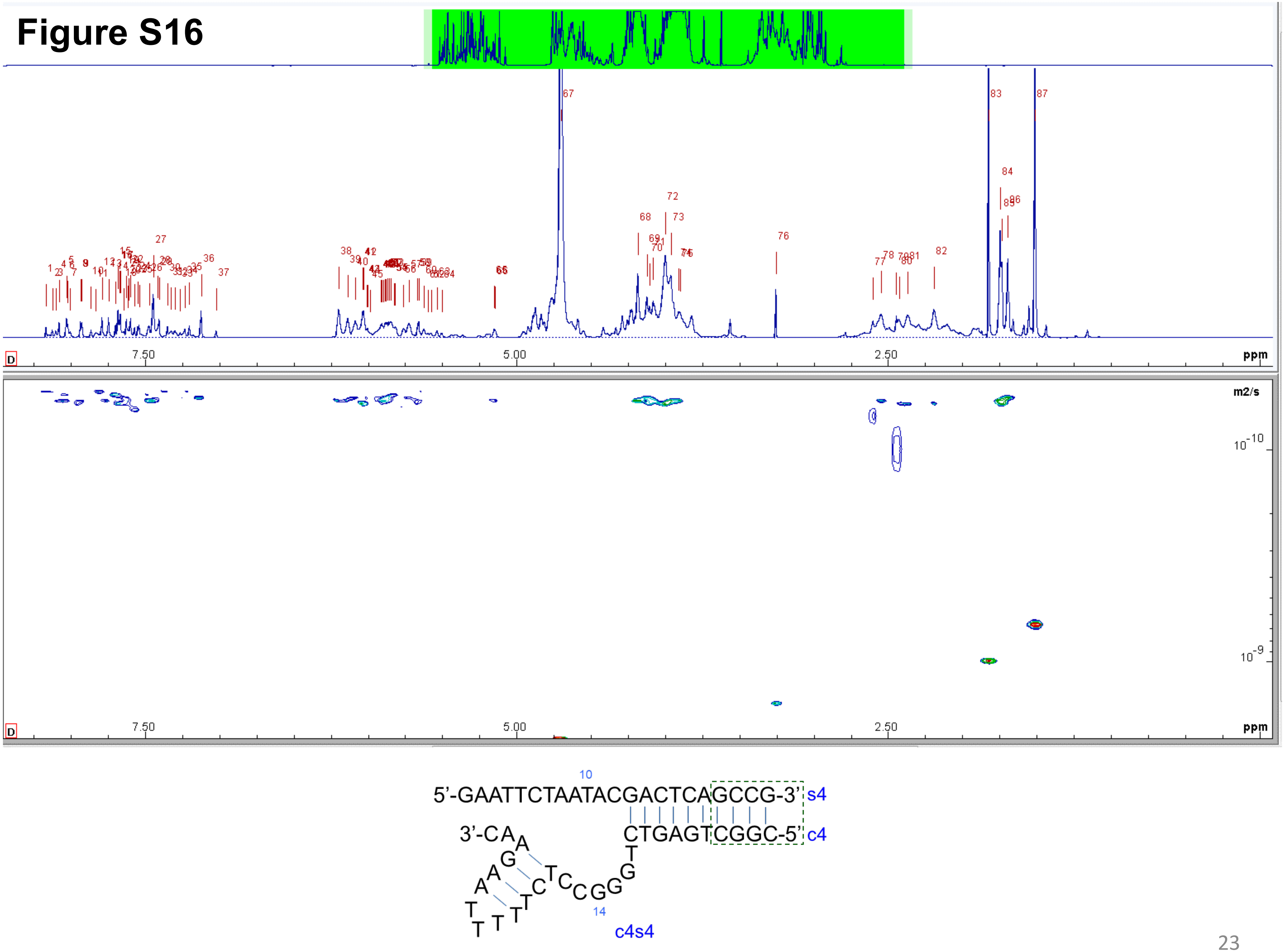

7.50
5.00
2.50
ppm
m2/s
10-10
10-9
10
5'-GAATTCTAATACGACTCAGCCG-3' s4
3'-CAA
CTGAGTCGGC-5' c4
14
c4s4

### Figure S17

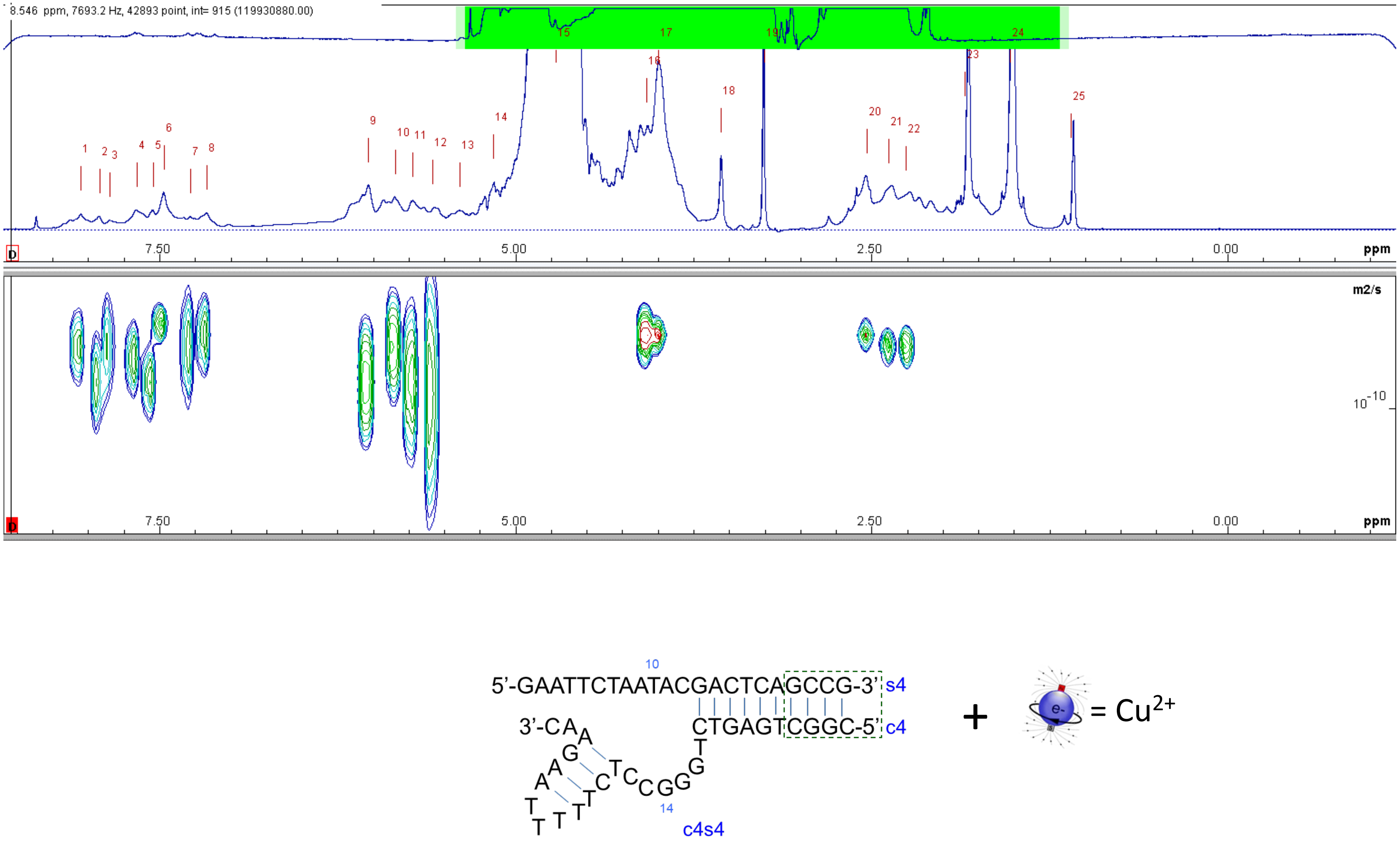
8.546 ppm, 7693.2 Hz, 42893 point, int= 915 (119930880.00)
1 2 3 4 5 6 7 8 9 10 11 12 13 14 15 16 17 18 19 20 21 22 23 24 25
7.50
5.00
2.50
0.00
ppm
m2/s
$10^{-10}$
10
5'-GAATTCTAATACGACTCAGCCG-3' s4
CTGAGTCGGC-5' c4
3'-CAA
14
c4s4
+
e-
= $Cu^{2+}$

#### Figure S18

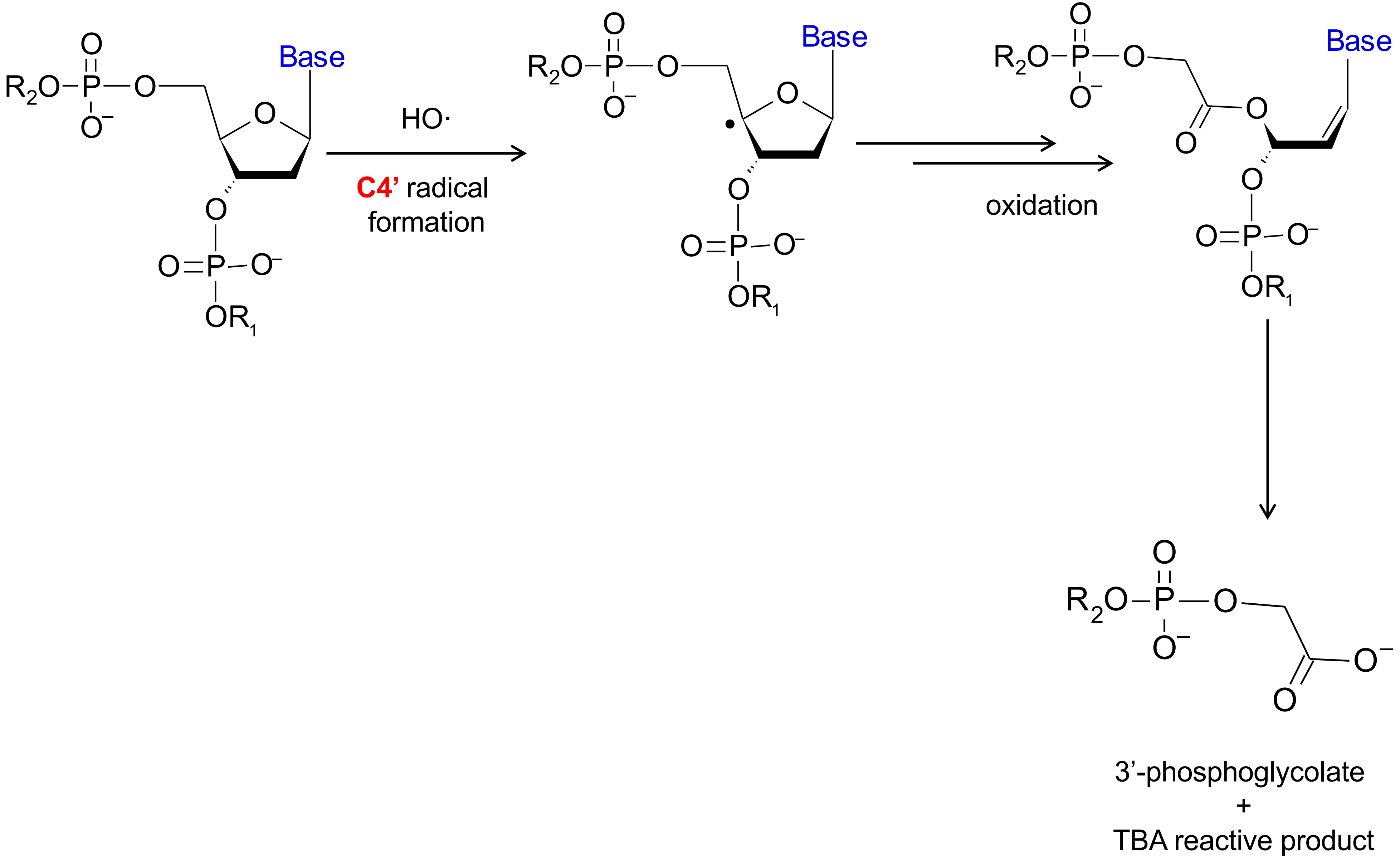

Base
HO·
C4' radical formation
oxidation
3'-phosphoglycolate
+
TBA reactive product

#### Figure S19

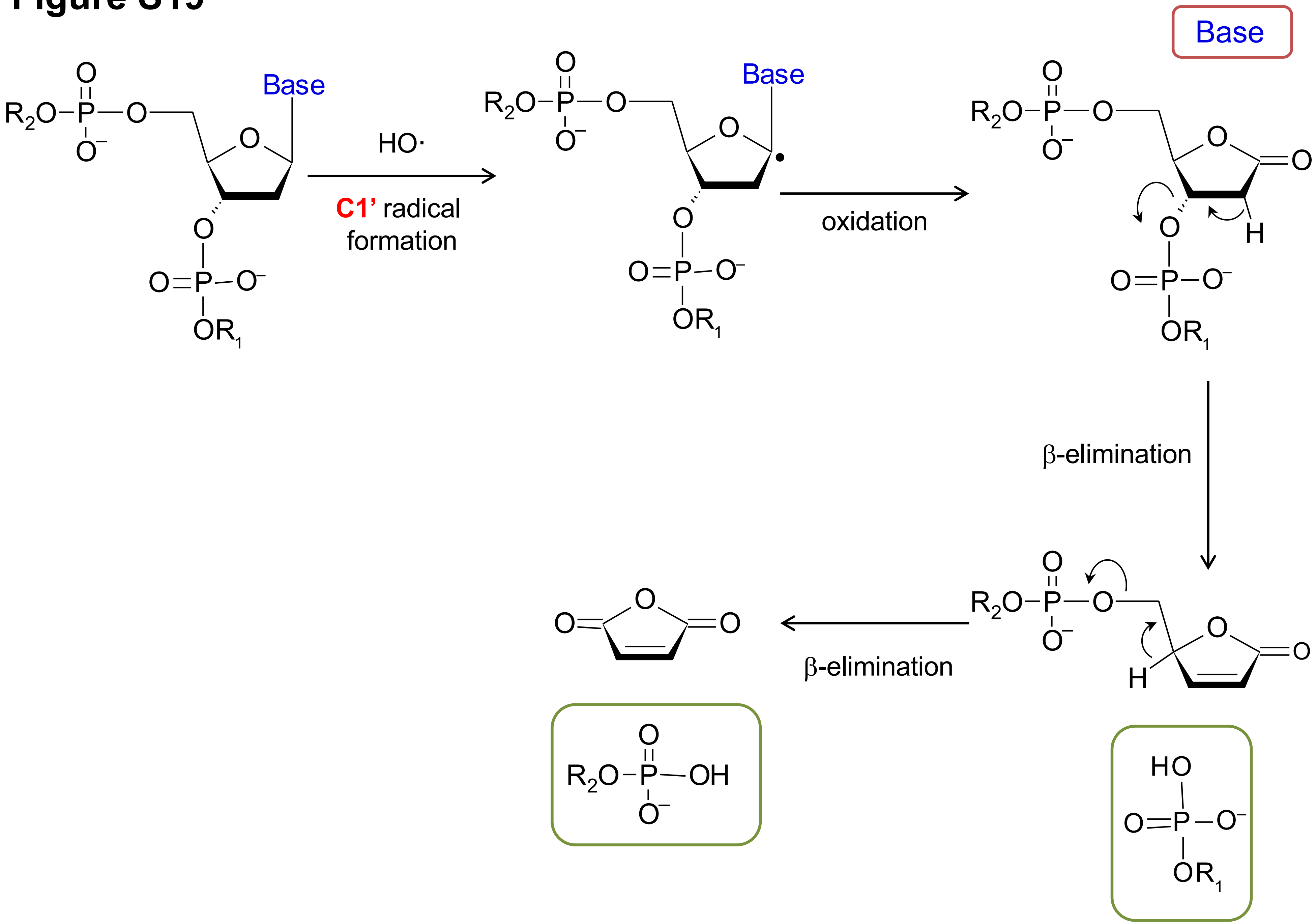
Base
HO·
C1' radical formation
oxidation
β-elimination
β-elimination

### Figure S20

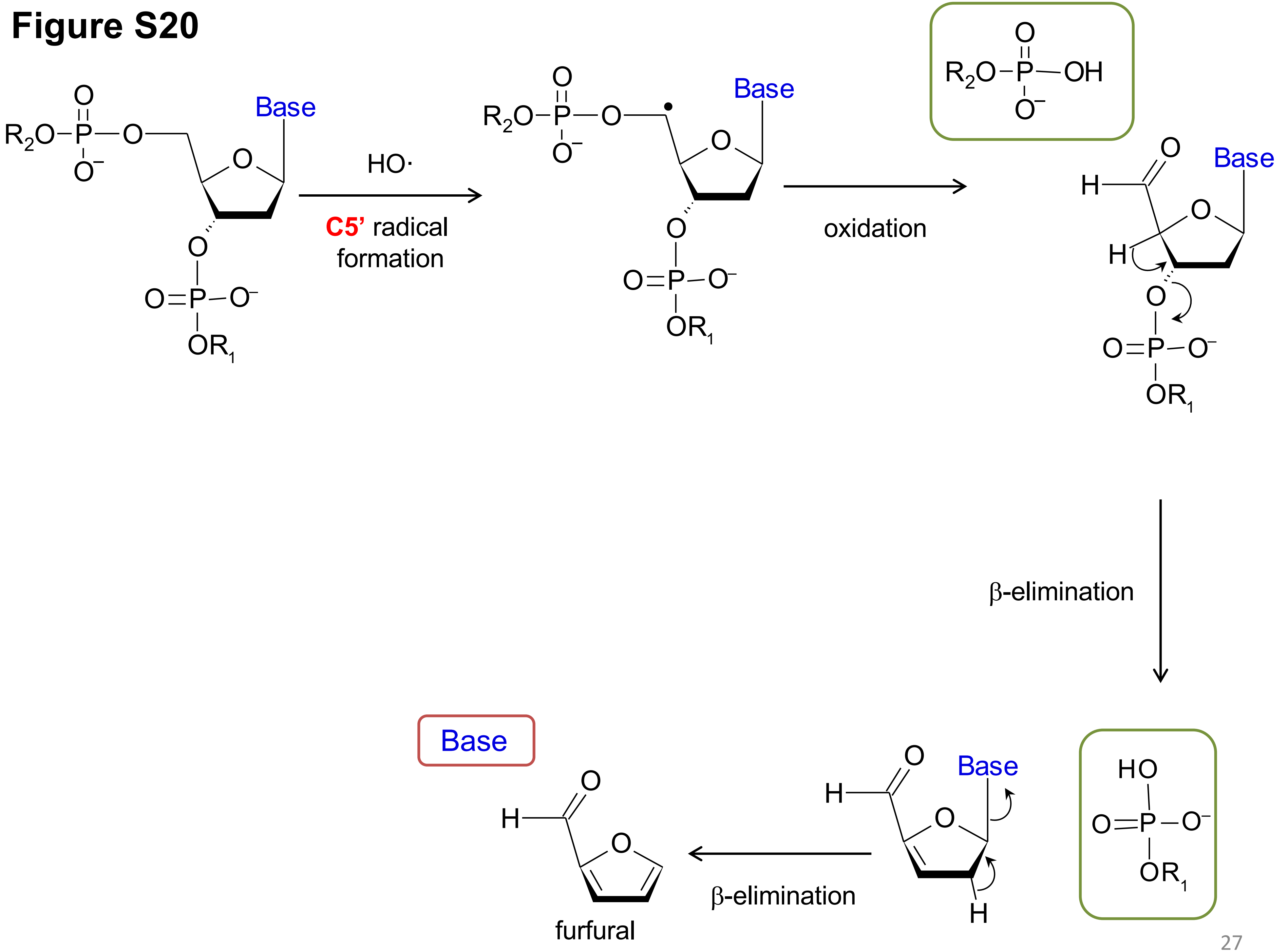

Base
HO·
C5' radical
formation
Base
oxidation
Base
β-elimination
Base
Base
β-elimination
furfural

### Figure S21

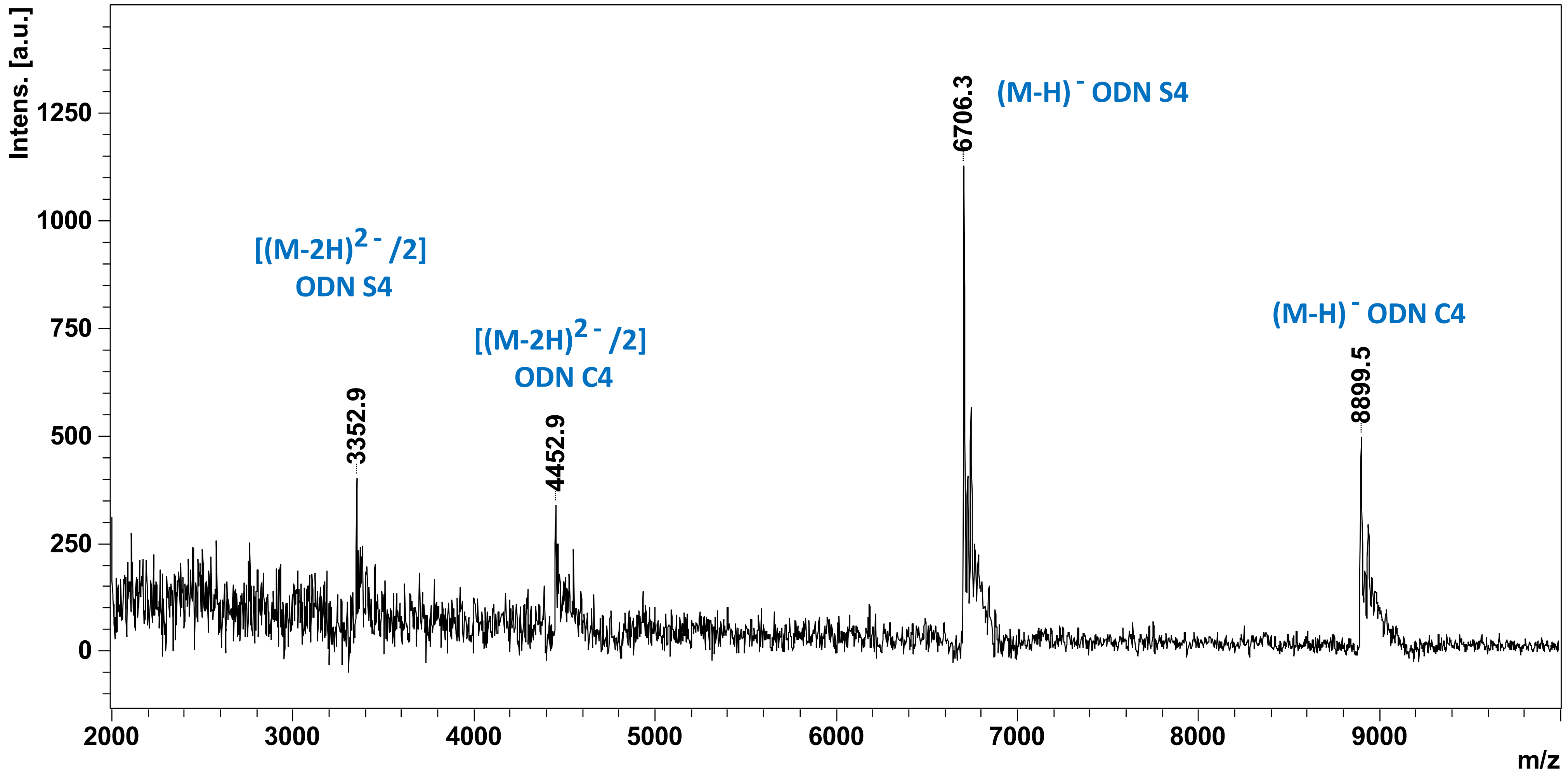

Intens. [a.u.]
1250
1000
750
500
250
0
2000
3000
4000
5000
6000
7000
8000
9000
m/z
[(M-2H)2 - /2]
ODN S4
3352.9
[(M-2H)2 - /2]
ODN C4
4452.9
6706.3
(M-H) - ODN S4
(M-H) - ODN C4
8899.5

### Figure S22

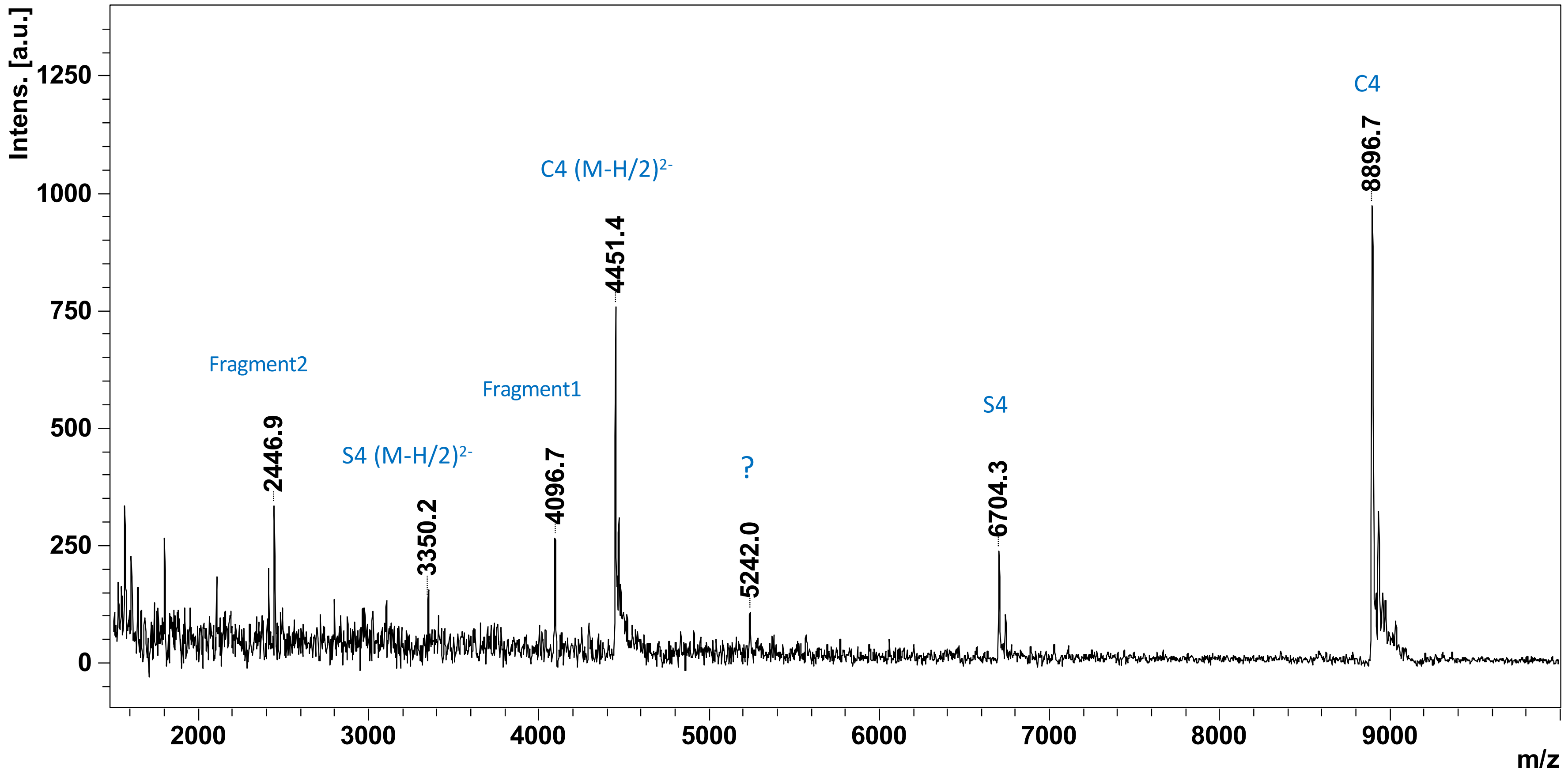

Intens. [a.u.]
1250
1000
750
500
250
0
C4
8896.7
C4 (M-H/2)2-
4451.4
Fragment2
2446.9
Fragment1
4096.7
S4 (M-H/2)2-
3350.2
?
5242.0
S4
6704.3
2000
3000
4000
5000
6000
7000
8000
9000
m/z

### Figure S23

5’-GGATTCTAATACGACTCAGCCG-3’ MW 6703,4

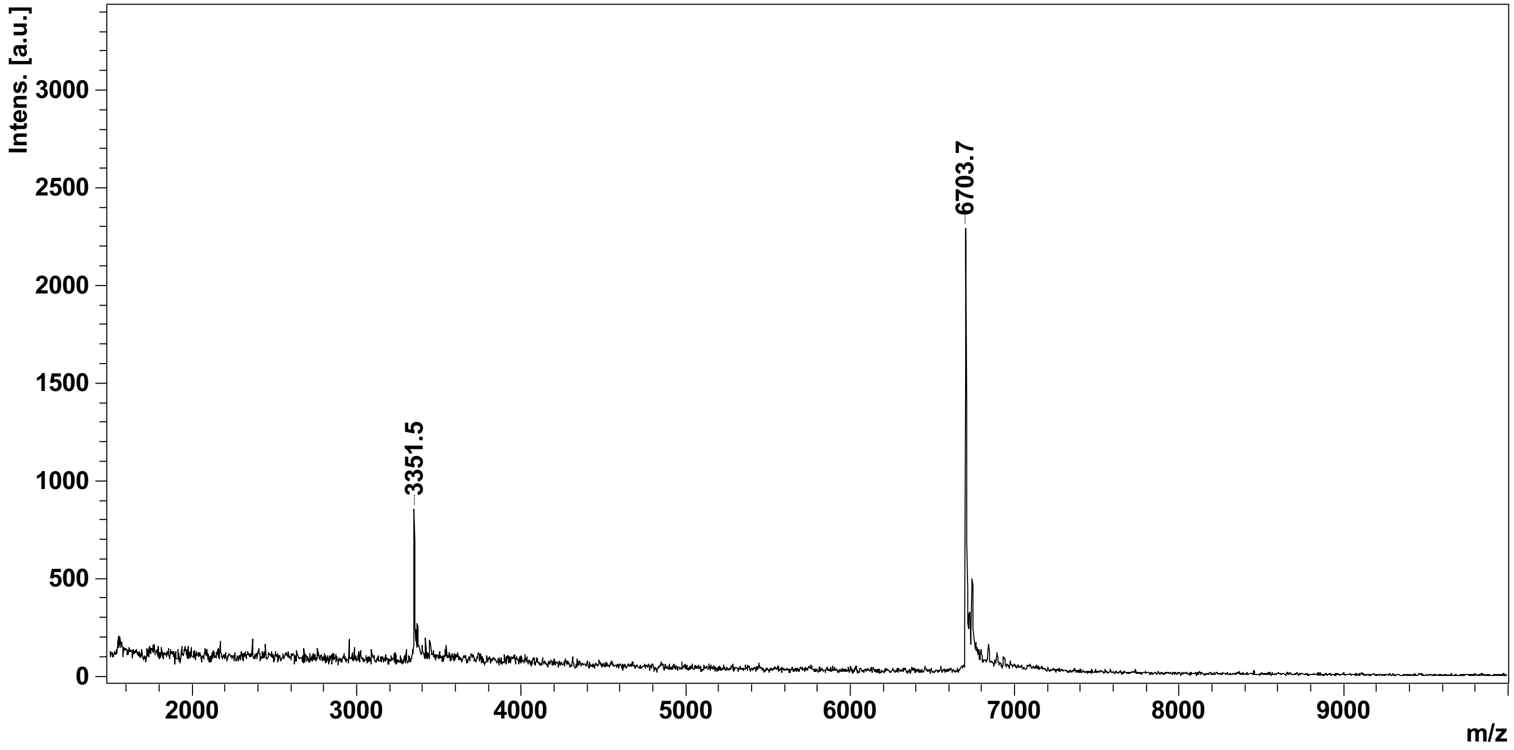

### Figure S24

5’-CGGCTGAGTCTGGGCCTCTTTTTAAGAAC-3’  MW 8899,8 Da

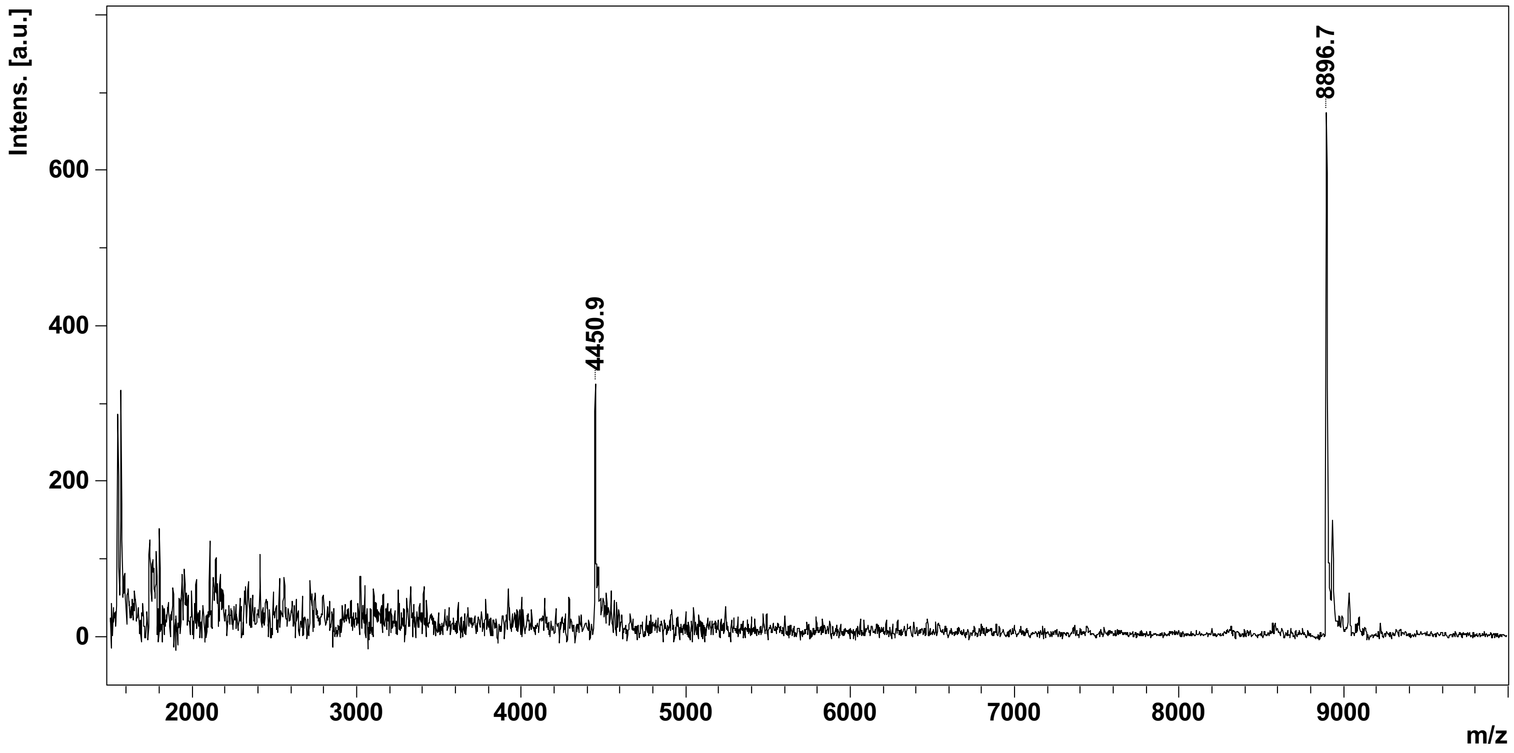

### Figure S25

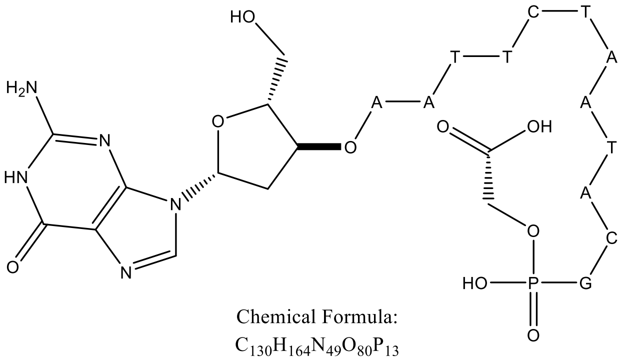


Chemical Formula:
$C_{130}H_{164}N_{49}O_{80}P_{13}$
Molecular Weight:
4095,6639

Fragment1 : ODN 3'phosphoglycolate

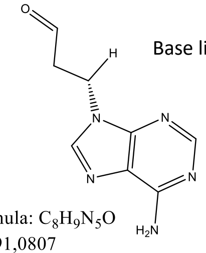


Base libérée

Chemical Formula: $C_8H_9N_5O$
Exact Mass: 191,0807
Molecular Weight: 191,1940

Chemical Formula: $C_{76}H_{99}N_{29}O_{49}P_8$
Molecular Weight: 2450,5721

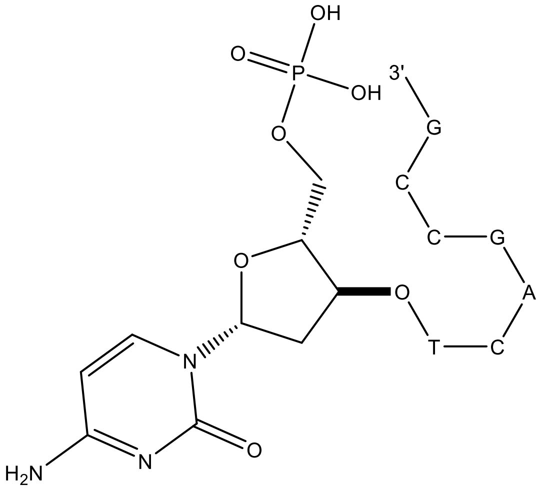


Fragment 2 : ODN 5' phosphate

### Figure S26

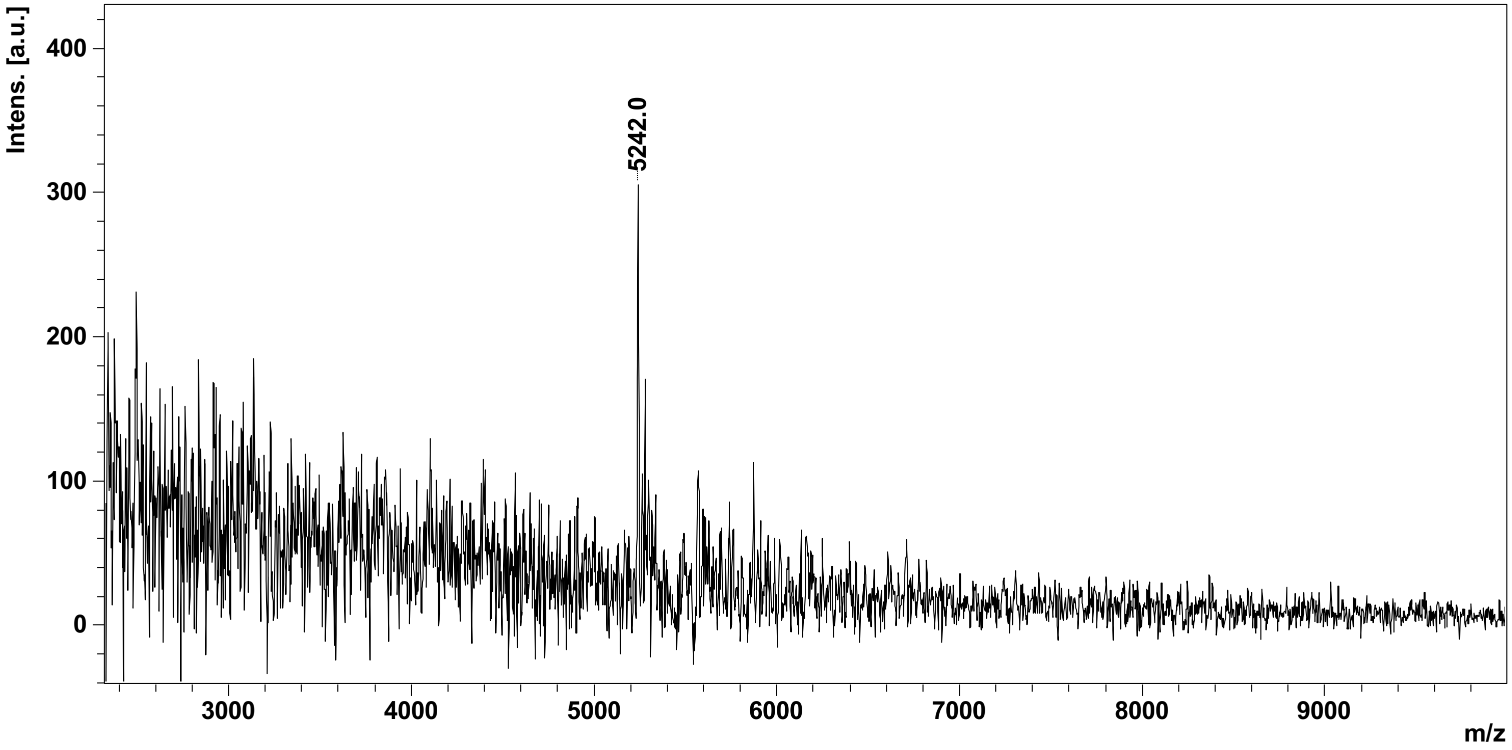
Intens. [a.u.]
400
300
200
100
0
5242.0
3000
4000
5000
6000
7000
8000
9000
m/z

### Figure S27

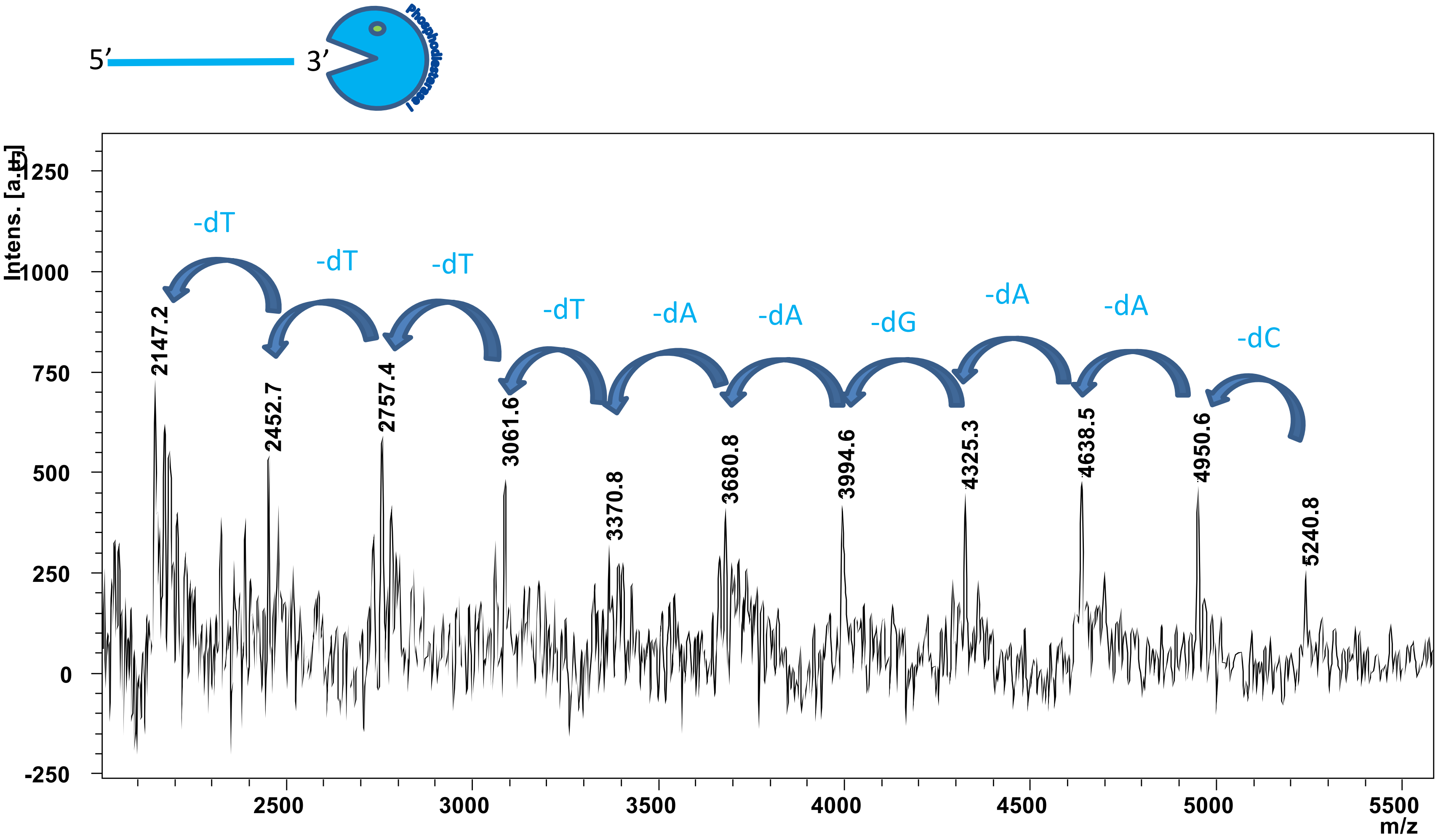

5'
3'
Phosphodiesterase I
-dT
-dT
-dT
-dT
-dA
-dA
-dG
-dA
-dA
-dC
2147.2
2452.7
2757.4
3061.6
3370.8
3680.8
3994.6
4325.3
4638.5
4950.6
5240.8
Intens. [a.u.]
1250
1000
750
500
250
0
-250
2500
3000
3500
4000
4500
5000
5500
m/z

### Figure S28

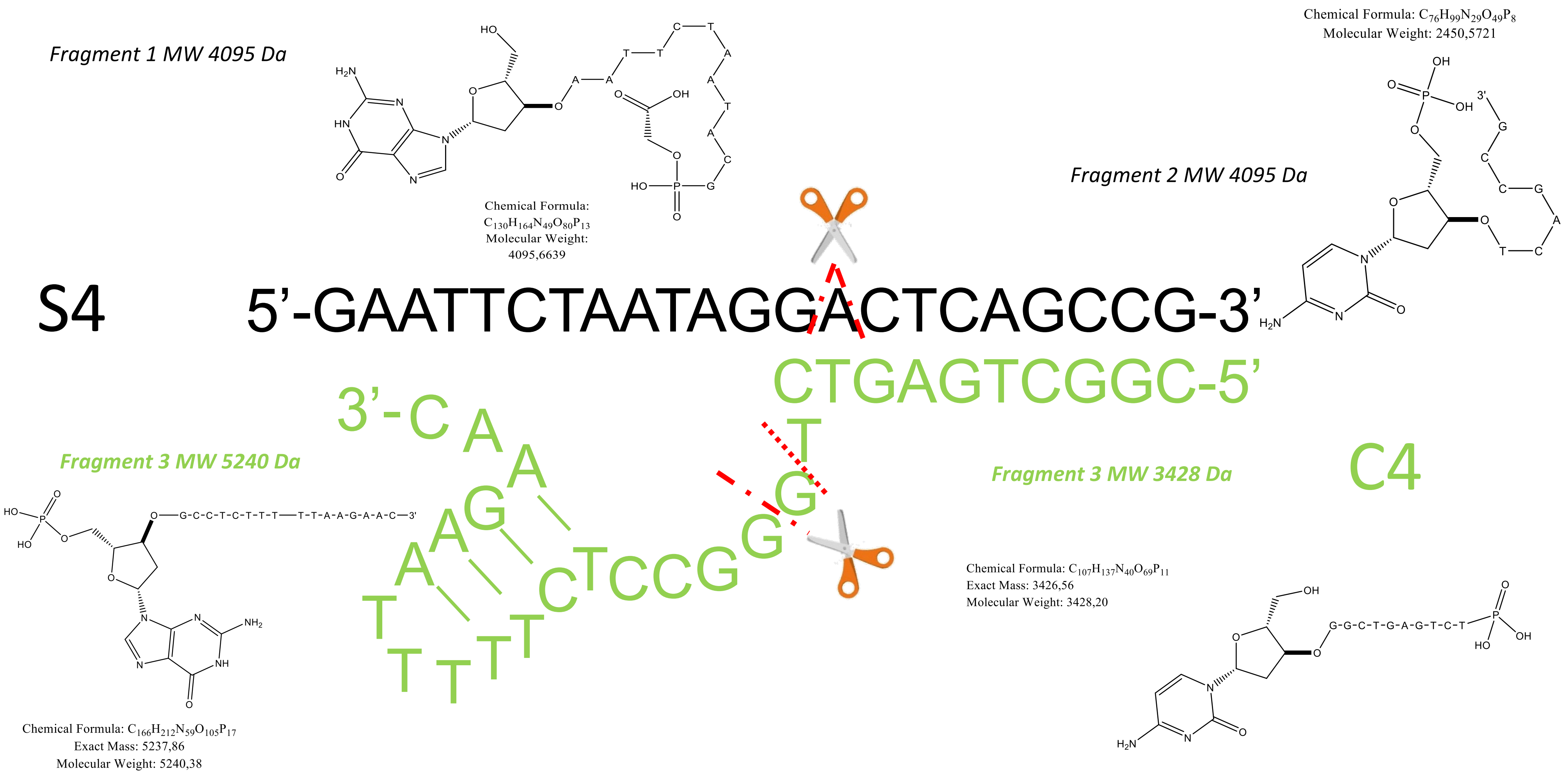

Fragment 1 MW 4095 Da
Chemical Formula: C130H164N49O80P13
Molecular Weight: 4095,6639
Chemical Formula: C76H99N29O49P8
Molecular Weight: 2450,5721
Fragment 2 MW 4095 Da
S4
5’-GAATTCTAATAGGACTCAGCCG-3’
CTGAGTCGGC-5’
3’-C
Fragment 3 MW 5240 Da
Fragment 3 MW 3428 Da
C4
Chemical Formula: C107H137N40O69P11
Exact Mass: 3426,56
Molecular Weight: 3428,20
Chemical Formula: C166H212N59O105P17
Exact Mass: 5237,86
Molecular Weight: 5240,38
