## Supplementary information list for "Cleaving DNA with DNA: cooperative tuning of structure and reactivity driven by copper ions"

[1]Department of Computer Science – Synthetic Biology Theme, Brunel University London, Kingston Lane, Uxbridge UB8 3PH, London, United Kingdom. [2]CNRS UMR 8516, Lille University, LASIRE, C4 Building, Avenue Paul Langevin, F–59655 Villeneuve d'Ascq, France. [3]Michel-Eugène Chevreul Institute (FR2638), Avenue Paul Langevin, F–59655 Villeneuve d'Ascq, France. [4]Grenoble University, UMR 5819 INAC–SyMMES-CREAB, CEA-CNRS-UGA, F–38000 Grenoble, France.

**Table of contents**

**S1.** Putative structures for 69-mer, 46-mer and c4s4 oligomers

**S2.** Modelled structure of 46-mer with Cu2+ randomly placed around the simulation box

**S3.** Cluster representatives and the respective population percentage of the cluster from the MD ensemble of 46-mer without $Cu^{2+}$

**S4.** Cluster representatives and the respective population percentage of the cluster from the MD ensemble of 46-mer with $Cu^{2+}$

**S5.** Distance distributions profiles obtained from MD ensembles with (MD-46mer-$Cu^{2+}$) and without $Cu^{2+}$ (MD-46mer) between the phosphate atoms of C6 and A21 and G19 and C46.

**S6.** Interaction between Cu2+ and the 46mer with a distance cut-off of 0.4nm was defined as residence time and the contact frequency as percentage of simulation time is shown for each independent MD trajectories below

**S7.** Time series distance profiles between the phosphate atoms of A14 and A41 calculated from the independent trajectories of MD simulations with Cu2+ (MD46mer- Cu2+) are shown below

**S8.** Spin labeled c4s4 oligomer and structures of nitroxide spin labels

**S9.** HYSCORE and HYSCORE fitting for the 15N guanosine on the catalyst c4 (G14)

**S10.** Table containing details of cross peaks for unlabeled and labeled (15N) c4s4

**S11.** Fitting (red line) for the DMPO-OH adduct

**S12.** Log(Molecular Weight) versus Log(Diffusion coefficient) obtained by the NMR experiments

**S13.** Table containing the details for the plot shown in S12

**S14.** NMR spectrum (900 MHz) monomeric guanosine

**S15.** NMR spectrum (900 MHz) monomeric guanosine with $Cu^{2+}$

**S16.** NMR spectrum (900 MHz) c4s4

**S17.** NMR spectrum (900 MHz) c4s4 with $Cu^{2+}$

**S18.** Putative mechanism on C4' hydrogen abstraction

**S19.** Putative mechanism on C1' hydrogen abstraction

**S20.** Putative mechanism on C5' hydrogen abstraction

**S21.** MALDI-TOF on the c4s4 double-strand

**S22.** MALDI-TOF on the cleaved c4s4

**S23.** MALDI-TOF control experiment on s4

**S24.** MALDI-TOF control experiment on c4

**S25.** Products of cleavages and corresponding molecular weight

**S26.** MALDI-TOF (zoom) on the peak 5240.8 m/z

**S27**. Scheme of enzymatic digestion on the fragment 5240.8 m/z

**S28.** Identification of the fragment 5240.8 m/z as product of catalyst decomposition
